# Gene Network Analyses Identify Co-regulated Transcription Factors and BACH1 as a Key Driver in Rheumatoid Arthritis Fibroblast-like Synoviocytes

**DOI:** 10.1101/2023.12.28.573506

**Authors:** Aurelien Pelissier, Teresina Laragione, Carolyn Harris, María Rodríguez Martínez, Percio S. Gulko

**Affiliations:** IBM Research Europe, 8803 Rü schlikon, Switzerland; Department of Biosystems Science and Engineering, ETH Zurich, 4058 Basel, Switzerland; Currently at Institute of Computational Life Sciences, ZHAW, 8400 Winterthur, Switzerland; Division of Rheumatology, Icahn School of Medicine at Mount Sinai, 10029 New York, United States; Currently at Yale School of Medicine, 06510 New Haven, United States

**Author notes:** T. Laragione contributed equally with A. Pelissier. M. R. Martinez contributed equally with P. S. Gulko.

## Abstract

RNA-sequencing and differential gene expression studies have significantly advanced our understanding of pathogenic pathways underlying Rheumatoid Arthritis (RA). Yet, little is known about cell-specific regulatory networks and their contributions to disease. In this study, we focused on fibroblast-like synoviocytes (FLS), a cell type central to disease pathogenesis and joint damage in RA. We used a strategy that computed sample-specific gene regulatory networks (GRNs) to compare network properties between RA and osteoarthritis FLS. We identified 28 transcription factors (TFs) as key regulators central to the signatures of RA FLS. Six of these TFs are new and have not been previously implicated in RA, and included BACH1, HLX, and TGIF1. Several of these TFs were found to be co-regulated, and BACH1 emerged as the most significant TF and regulator. The main BACH1 targets included those implicated in fatty acid metabolism and ferroptosis. The discovery of BACH1 was validated in experiments with RA FLS. Knockdown of BACH1 in RA FLS significantly affected the gene expression signatures, reduced cell adhesion and mobility, interfered with the formation of thick actin fibers, and prevented the polarized formation of lamellipodia, all required for the RA destructive behavior of FLS. This is the first time that BACH1 is shown to have a central role in the regulation of FLS phenotypes, and gene expression signatures, as well as in ferroptosis and fatty acid metabolism. These new discoveries have the potential to become new targets for treatments aimed at selectively targeting the RA FLS.

## Introduction

Rheumatoid arthritis (RA) is a common systemic autoimmune and inflammatory disorder characterized by synovial inflammation and hyperplasia that may lead to joint destruction[1, 2, 3]. New biologic disease-modifying anti-rheumatic drugs (bDMARDs) and JAK inhibitors that target various inflammatory pathways have significantly improved disease control and outcomes [4, 5]. Yet, a considerable number of RA patients still have only partial or no response to therapy, and sustained remission remains uncommon [6]. The development and progression of RA involves dynamic interactions between multiple genetic and environmental factors, and therefore, understanding the heterogeneous pathophysiological processes in RA patients remains a major challenge [7]. Among the cell types found in synovial tissues, fibroblast-like synoviocytes (FLS) [8, 9] are centrally relevant in the RA pathogenesis [8, 10, 11]. In RA, FLS become activated producing increased levels of inflammatory mediators that contribute to driving the local inflammatory response and joint damage [12]. FLS also have increased invasive properties that together with their increased production of proteases contribute to cartilage and bone damage [8]. While past studies have linked the invasiveness of FLS with the severity of joint damage, both in RA patients [13] and rodent models [14], the underlying mechanisms of those processes remain incompletely understood.

Genome-wide association studies (GWAS) and differential gene expression (DEG) studies have significantly improved our understanding of the disease’s genetic underpinnings [15, 16]. Yet, those studies did not differentiate the contribution of individual cell types to disease. In this context, mapping the identified transcriptional and immune signatures specific to FLS could substantially expand our understanding of RA disease processes[17]. Identifying transcription factors (TFs) and their associated regulatory signatures is of particular interest, as TFs play a pivotal role in regulating gene expression [18]. Moreover, with an estimated count of 1,000 TFs in humans [19], identifying those that govern the phenotypic traits of FLS in RA may open new possibilities for novel therapeutic target discovery. Recently, single-cell RNA sequencing studies are offering valuable insights into understanding diseases at a cell-specific level [20, 21]. Still, utilizing these datasets presents several challenges due to limited patient numbers, batch effects, and sparse data [22]. Consequently, inferring networks based on these datasets has proven to be particularly difficult [23, 24, 25].

In this study, we provide a comprehensive analysis of gene regulation in RA FLS. We leverage FLS-specific gene expression data [20] to construct sample-specific gene regulatory networks (GRNs) in both RA and controls. Unlike previous studies in RA relying on the inference of a cohort-specific GRN [26, 27, 7], our approach allowed us to gain new insights by characterizing the disease through differential analysis of the sample-specific network edges, from which we identified and implicated specific TFs in driving FLS-specific expression differences. We identified several new key regulators in RA and identified BTB and CNC Homology 1 (BACH1) and H2.0 like homeobox (HLX) among the most significant and specific regulators in FLS. To validate our discovery strategy, we conducted experiments silencing BACH1 in FLS cell lines from RA patients, demonstrating its influence on FLS migration, adhesion, and lamellipodia formation. Our study is an important step towards the development of novel RA treatments with FLS-specific pathway targeting.

## 1 Results

### 1.1 Differentially expressed genes and pathways in RA versus OA

We used cell type-specific RNA-seq data from FLS [20] to obtain gene expression profile for 18RA and 12 OA biopsies. Like in most RA FLS studies, OA samples were used as controls given the lack of available normal biopsies. Next, we computed the differential gene expression in each dataset (Student’s *t*-test) between RA and OA samples. This means that for each gene, we obtain a differential expression score (denoted *t*_diff-expr_) from which we can extract a set of DEGs (Supplementary Table S1). The genes with the most significant difference included the long intergenic non-protein coding RNA 1600 (LINC01600), the solute carrier family 25 member 26 (SLC25A26), and monocyte to macrophage differentiation-associated (MMD; Figure 1A). Pathway analyses were done including 1,093 genes with a *t*_diff-expr_ *>* 2, and revealed an overrepresentation of “negative regulation of low-density lipoprotein receptor activity”, followed by others such as axon guidance and carnitine shuttle (Figure 1B, Supplementary Table S2).

**Figure 1:**
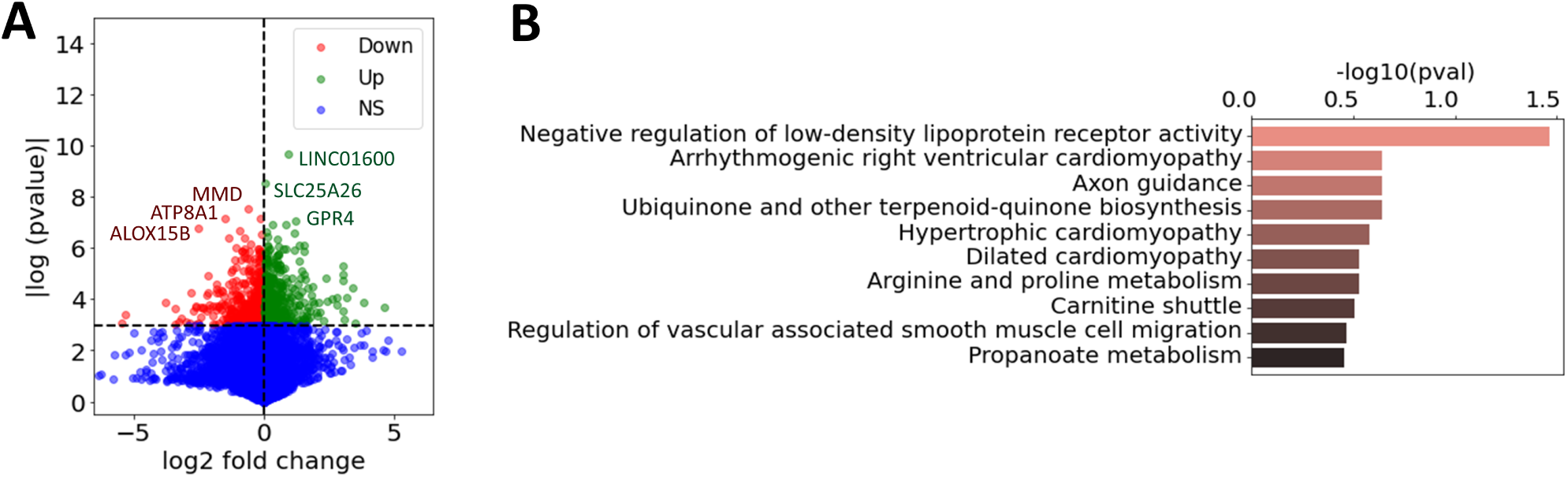
(A) Volcano plot of the computed log2 fold change and *p*-values for all the expressed genes in RA *vs* OA FLS (Student’s *t*-test), where the six genes with the highest *p*-values were marked. Genes were labeled as downregulated, upregulated, or non-significant (NS). (B) Most enriched pathways of the differentially expressed genes in RA vs OA FLS (1093 genes with *|t*_diff-expr_*| >* 2), combined from GO [28], KEGG [29] and Reactome [30], ranked by adjusted p-values.

### 1.2 Key TF regulators and pathways driving regulatory differences between RA and OA FLS

Gene expression phenotypes are more effectively interpreted through the identification of the TFs that regulate them. Therefore, we next identified the TFs regulating RA phenotypes by inferring sample-specific GRNs and identifying differential regulatory patterns between RA and OA. We aimed to get insight into the regulatory relationships between DEGs, and whether they involved TF-mediated inhibition or activation, and co-regulations. To explore this question, we constructed FLS-specific gene regulatory networks by integrating RA gene expression datasets [20] with prior knowledge about TF binding motifs (from the Catalog of Inferred Sequence Binding Preferences, CIS-BP [31]) and about protein-protein interactions (from StringDB [32]). Briefly, we leveraged LIONESS [33] to compute individual gene regulatory networks for each biopsy sample (Methods 2.3, Figure 2A). These networks incorporate edge weights to quantify the likelihood of regulatory interactions between TFs and their target genes (TGs). We then compared these edge weights between RA and OA samples using a Student’s t-test and obtained a score *t*_diff-edge_ for each edge TF *→* TG. From these edge-specific differential analyses, we generated a novel network, the *differential GRN* (dGRN), which illustrates the differential regulation between RA and OA (Figure 2B). To quantify TFs’ regulatory function, we calculated a TF regulatory score (*t*_reg_). This score is calculated based on the average absolute differential weight of the regulatory edges between each TF and its associated TGs, i.e. average of |*t*_diff-edge_| between a TF and its targets (Methods 2.4).

**Figure 2:**
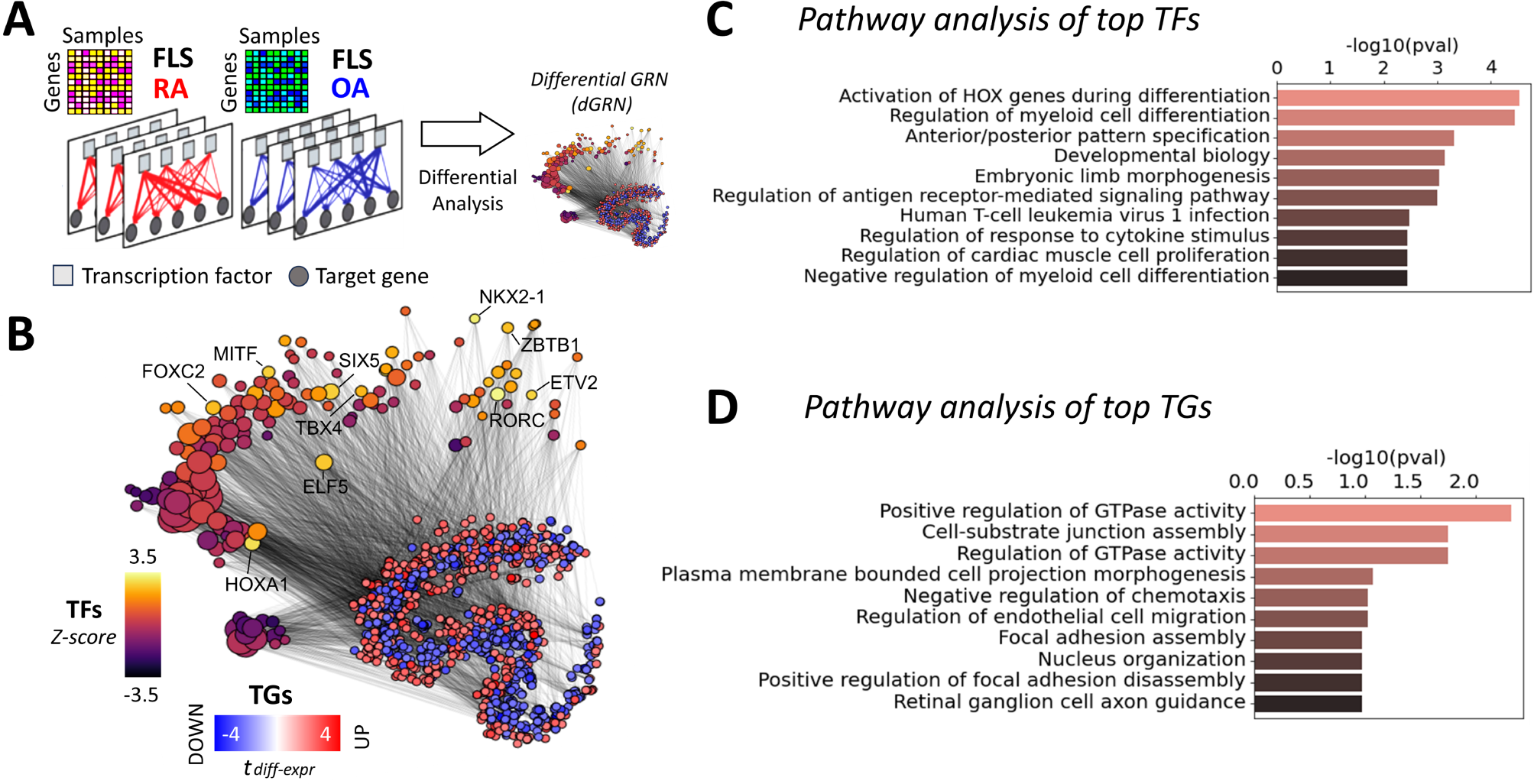
(A) 18 RA and 12 OA FLS networks were inferred from FLS gene expression profiles [20]. Networks incorporate edge weights representing the probability of regulatory interactions between transcription factors (TFs) and their target genes (TGs). The differential analysis of the edge weights between RA and OA resulted in the assembly of a *differential GRN* (dGRN) specific to RA FLS. (B) t-SNE representation of the dGRN of RA FLS. For visual clarity, only TGs with a differential expression score in FLS above 2 (*t*_diff-expr_ > 2), and only edges with a differential edge weight score above 2 (*t*_diff-edge_ > 2), are shown. TF node sizes are proportional to the node degree, and colored according to their regulatory score in the network. Gene nodes are colored according to their differential expression score in FLS. The top 10 TFs ranked by their regulatory scores are labeled (C) Top 10 enriched pathways of the major TFs involved in RA FLS, compiled from GO [28], KEGG [29] and Reactome [30], and ranked by their adjusted *p*-values. (D) Top 10 enriched pathways of the major TGs involved in RA FLS, ranked by adjusted *p*-values.

We identified 185 TF signatures in RA FLS, and show the top 20 ranked by their regulatory scores (Table 1, with Z-statistics of each TF in parentheses). The Z-statistics quantifies the number of standard deviations by which the score of a given TF deviates from the mean score across all TFs. For additional insight into the important pathways involved in RA FLS, we selected the 185 TFs with a Z-statistics above 0.5 (Supplementary Table S3). This collection constitutes our set of key *TF drivers* in RA FLS. Using this set, we performed a pathway enrichment analysis using the Gene Ontology database (GO) [28], KEGG [29] and Reactome [30] databases. Given that all TFs are DNA-binding proteins, we removed any terms associated with RNA and DNA transcription, as these processes are ubiquitous and therefore not likely to be specific to RA. After this filtering, 77 significantly enriched (adjusted *p*-value < 0.05) pathways were identified. The 20 most significant pathways included HOX genes activation, regulation of myeloid cell differentiation, and TNF signaling among others (Figure 2C, Supplementary Table S4).

**Table 1:** Top 20 ranked TF regulator in FLS with their Z-statistics provided in parentheses.

| Rank 1-5 | Rank 6-10 | Rank 11-15 | Rank 15-20 |
| --- | --- | --- | --- |
| RORC (3.39) | SIX5 (2.61) | CBFB (2.16) | ETV3 (2.04) |
| NKX2-1 (3.37) | ELF5 (2.46) | BBX (2.16) | TLX3 (1.97) |
| HOXA1 (3.20) | ZBTB1 (2.42) | TAL1 (2.13) | HOXB3 (1.95) |
| ETV2 (2.67) | FOXC2 (2.27) | ZBTB3 (2.11) | PKNOX1 (1.95) |
| MITF (2.64) | TBX4 (2.23) | RARG (2.09) | MECOM (1.94) |

In addition to evaluating the TF regulation of RA, we also investigated TGs that were differentially targeted by more than one the key TF drivers we identified (185 TFs with Z-statistics above 0.5). Interestingly, we found that some TGs were differentially targeted (*|t*_diff-edge_*| >* 1) by more than 30% of these TFs (the top targeted genes were ALDH1A2, SYNE2, and NF1). We provide the top 100 targeted genes in the Supplementary Table S5. A pathway enrichment analysis of the 211 TGs targeted by more than 20% of the key TF drivers in RA highlighted the pathways “Positive regulation of GTPase activity”, “Cell-substrate junction assembly” and “negative regulation of chemotaxis”, among others (Figure 2D, Supplementary Table S6).

### 1.3 An independent approach increases the confidence in the identified regulators

To enhance the reliability of the identified TF regulators and dGRN and mitigate variability commonly linked with reconstructed networks [34], we incorporated a set of 14 publicly available networks, sourced from the literature, which also incorporated edge weights as a measure of the interaction confidence between nodes in various cell types and tissues. Since there are no publicly accessible networks of synovial tissues or FLS, we selected networks derived from immune-related tissues, such as lymph nodes, spleen, tonsils, and blood, as well as various immune cell types (see Supplementary Table S8).

We used an approach based on the *Key Driver Analysis* (KDA) [35], a computational pipeline to uncover major disease-associated regulators or causative hubs in a biological network (Methods 2.6). In brief, using a list of RA-associated signatures as a starting point, we identified all potential *key drivers genes* (KDGs) linked to these signatures through network edges. From this set, genes with a higher number of connections to RA-associated genes than expected by chance were considered potential drivers. To increase the robustness of this approach, we performed a KDA based on a previously computational method [36] and using two independent lists of RA-associated gene signatures, including a list of DEGs and a list obtained from the literature (Supplementary Table S7). Combining the results from the two lists, we obtained a list of 174 key TF drivers that were identified in at least one of the 14 networks (Supplementary Table S9).

A list of potential key TF regulators in FLS was generated by combining our dGRN with the KDA approach (Table 2). Namely, we retained TFs (i) whose regulatory scores (*t*_reg_) computed with the dGRN network exceeded half a standard deviation above the mean score of all TFs (i.e. Z-statistics > 0.5), and (ii) were identified as a key drivers (KDG) by the KDA approach described above. We identified 28 TFs that met these two conditions. Among them, 7 were new TF regulators (HLX, BACH1, ETV7, TGIF1, ELF1, HIVEP1, and PLAGL1) not previously implicated in RA, and two of these (BACH1 and HLX) were specific to FLS, while the others were also associated with T and/or B cell signatures. The remaining TFs had been previously reported in RA studies, further validating our discovery strategy (Table 2).

**Table 2:**
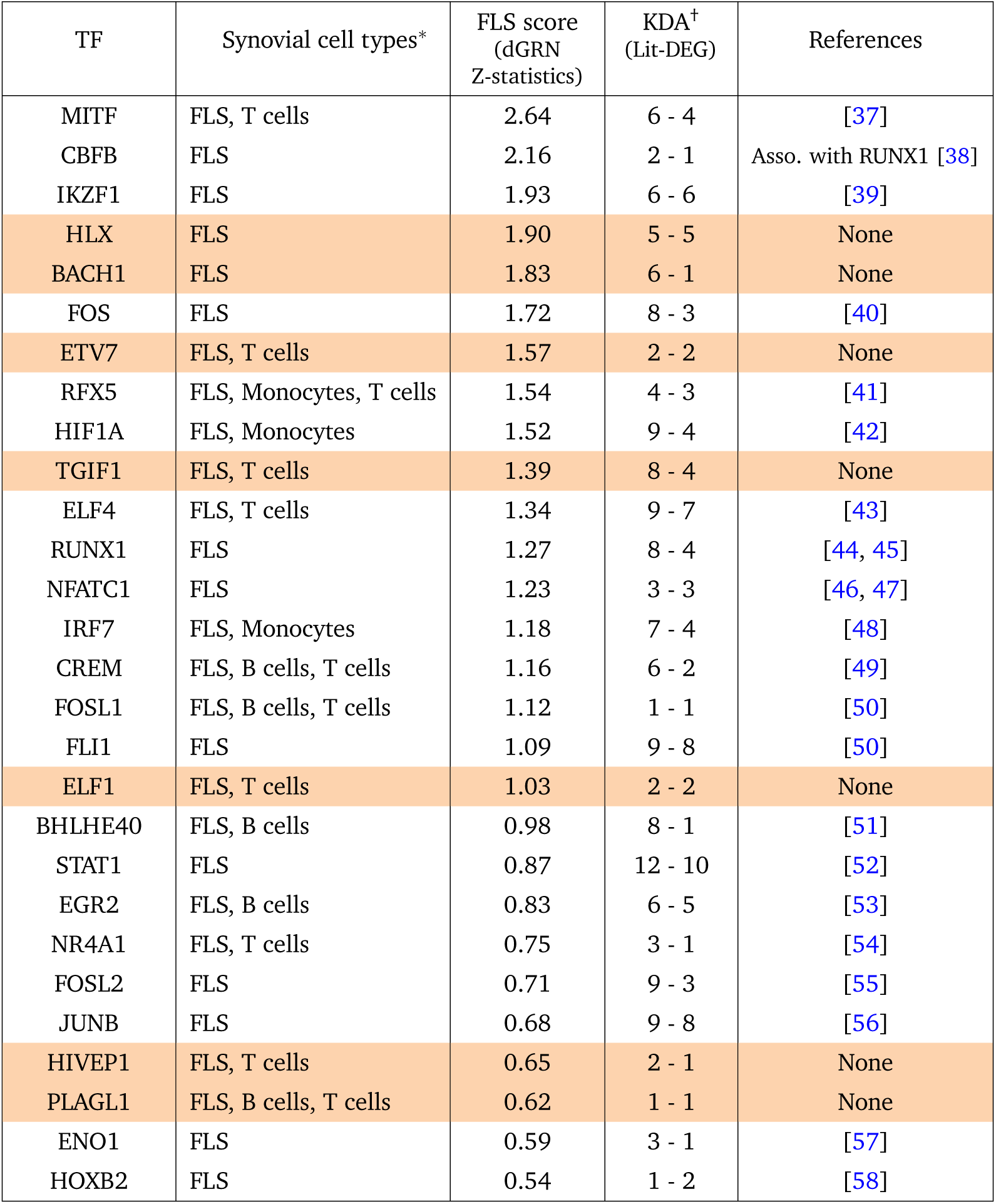
The 28 key TFs implicated in the regulation of FLS in RA, as identified in our analyses, are ranked by Z-statistics. A higher Z-statistic indicates a greater regulatory implication in FLS signatures. (*) Some of the key TFs in FLS were also found as key regulators in other synovial cell types [36]. (†) The left and right numbers correspond to the number of networks, among 14, in which the TF was identified as a KDG using, respectively, the literature and DEG list [36].

Importantly, the TFs shown in Table 1 are quite different from those on Table 2. Incorporating condition (ii) reduced our previous list from 185 TFs to 28 TFs. With this strategy, we increased the confidence in our results, as the chances of selecting false positives was reduced by combining the two criteria.

### 1.4 TF-TF co-regulation network in FLS

Having identified 28 significant TFs associated with RA in FLS, we next evaluated their co-regulatory activity, potentially revealing distinct clusters of co-regulation. TF-TF co-regulation was quantified using the Pearson correlation between the regulatory differential activities of the TFs and their TGs, i.e. *corr*(*t*_diff-edge_(*TF_i_*), *t*_diff-edge_(*TF_j_*)), focusing only on the TGs that are common to both TFs (Methods 2.5). We used a hierarchical clustering algorithm to identify groups of co-regulatory TFs using a correlation threshold of 0.5. This analysis identified 5 TF clusters, with four small and one major group containing 17 TFs (Figure 3A). This suggests a major co-regulated hub in RA FLS, with BACH1, HIF1A, TGIF1, and FOSL1 having a central key driving role (highest Z-statistics among TFs in the largest cluster). Among these, BACH1 has the highest Z-statistics (Supplementary Table S10, Figure 3A). These clusters were not completely independent, as there was also co-regulatory activity across TF belonging to different clusters (Figure 3B).

**Figure 3:**
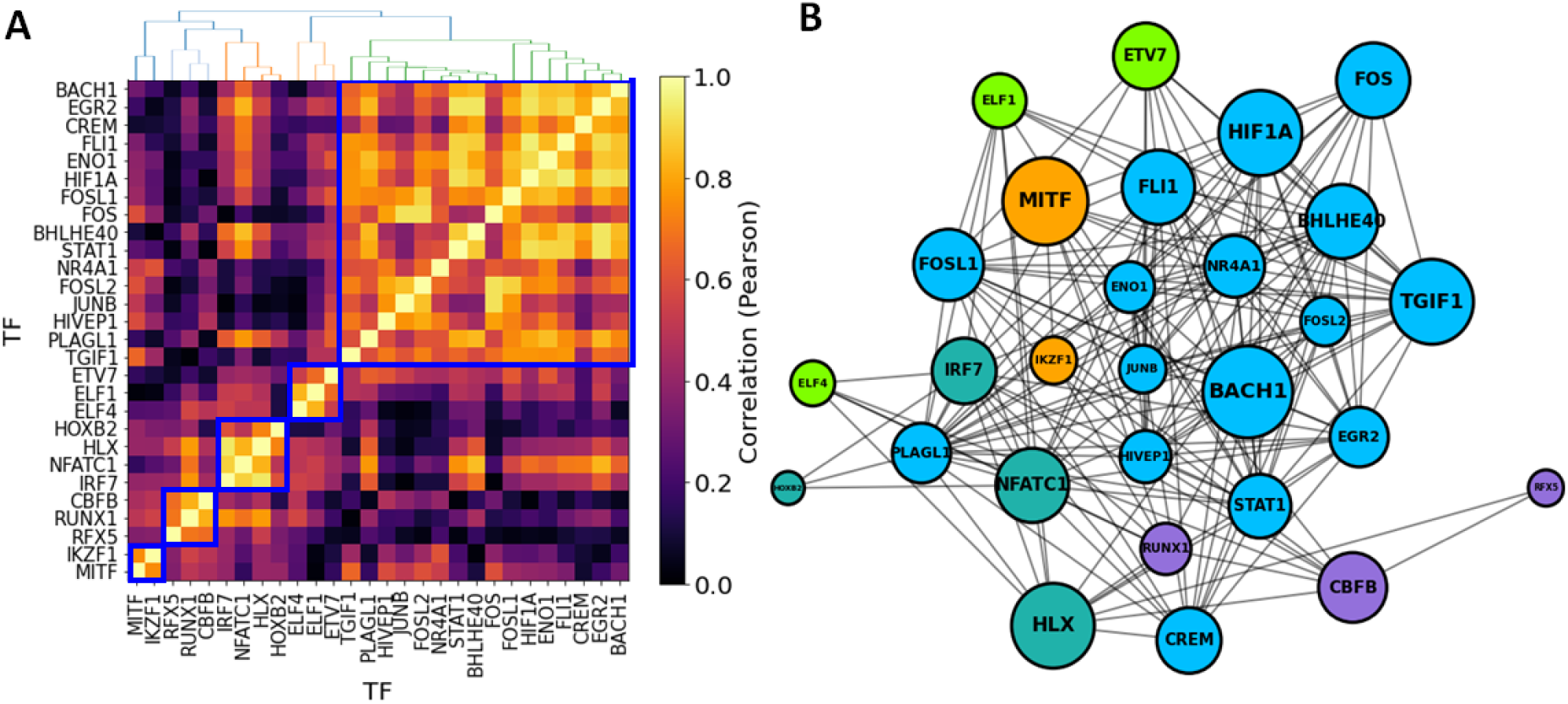
Co-regulatory activity of significant TFs in RA FLS. (A) Pairwise TF-TF co-regulation heatmap, quantified in terms of the Pearson correlation between the differential edge weight *t*_diff-edge_ to their common TGs (Methods 2.5). A hierarchical clustering approach was used to group the TFs into clusters (depicted in blue square). (B) Network visualization of the major TFs involved in FLS regulation in RA. Edges represent a correlation > 0.5, and node sizes are proportional to both the degree and the TF regulatory scores (*t*_reg_). Nodes are colored according to their cluster assignment.

### 1.5 BACH1 regulatory network in RA FLS

BACH1 was among the strongest TF regulators identified in RA FLS. To examine the regulatory effect of BACH1 on FLS, we constructed the network of the BACH1-target genes that were also DEGs between RA and OA (Figure 4A). This network included other significant TFs, such as PAX8, CBFB, NFE2L2, ETS1, RUNX1, and SMAD4, which also contributed to the co-regulation of major BACH1 targets. Besides these TFs, this network also included 131 genes that were differentially targeted by BACH1 (Supplementary Table S11). A pathway enrichment analysis of these genes showed an over-representation of the “fatty acid degradation” and “ferroptosis” pathways, driven by genes HADHB, ACSL6 and CPT1C among others (Figure 4B and Supplementary Figure S12). These findings are in line with the literature, where it has been reported that BACH1 promotes ferroptosis [59], and suppresses fatty acid biosynthesis [60]. These observations implicate for the first time BACH1 in the regulation of RA FLS metabolism and ferroptosis.

**Figure 4:**
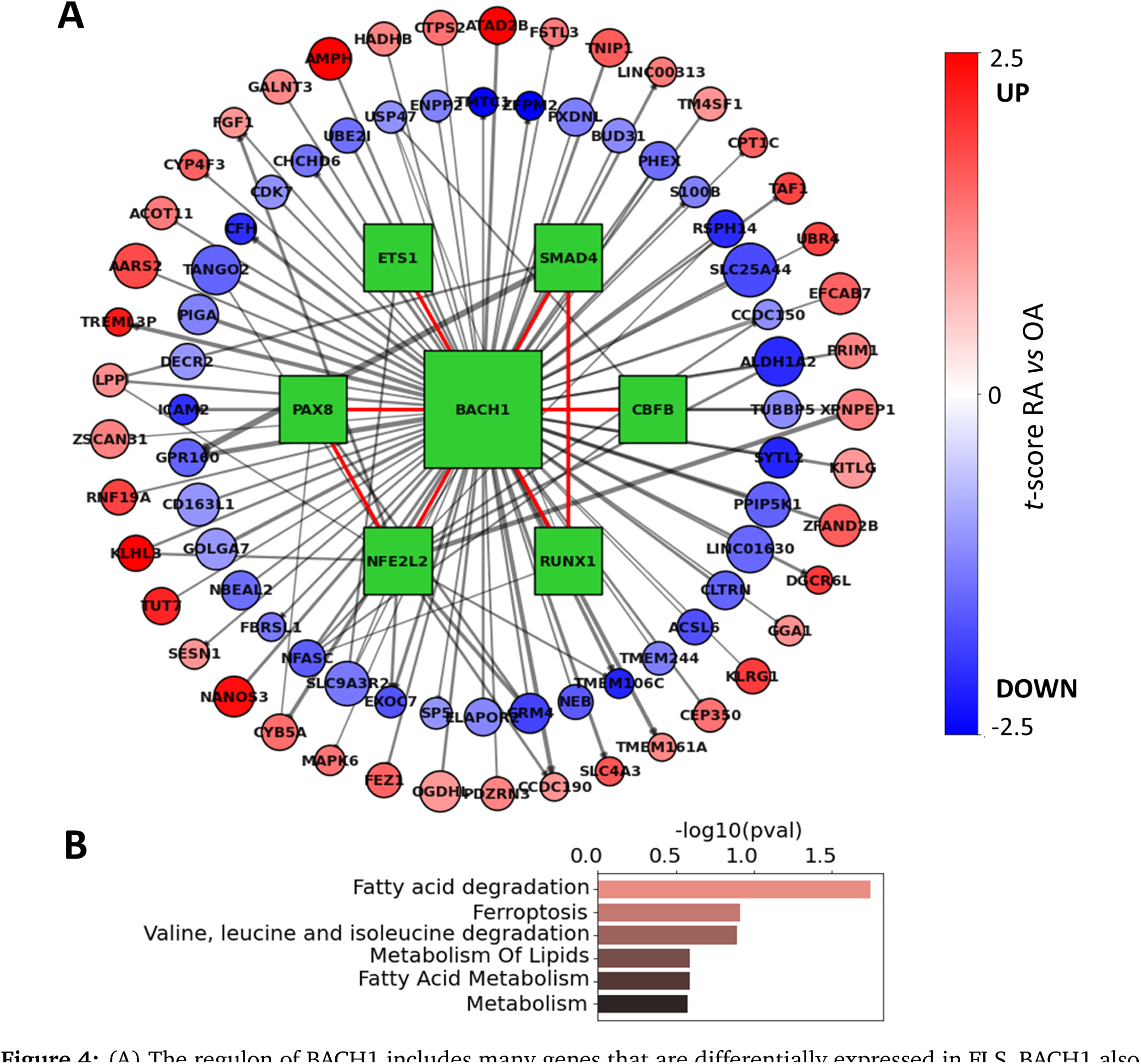
(A) The regulon of BACH1 includes many genes that are differentially expressed in FLS. BACH1 also interacts with 6 TFs (green squares), and in a tight cluster that contributes to the co-regulation of BACH1 target genes. Interactions between TFs are shown with a red edge, while TF-TG interactions are shown with a grey edge. Edge thickness is proportional to edge weights, node sizes are proportional to RA differential t-scores (*|t*_diff-edge_*|*), and target nodes are colored according to their RA differential expression in FLS (*t*_diff-expr_). (B) Top 5 over-represented pathways identified among the BACH1 differentially expressed target genes.

### 1.6 Effect of BACH1 knockdown on gene expression

To gain more direct insight into the role of BACH1 in regulating RA FLS, we ranked the BACH1 regulatory weights (average likelihood of TF-TG interaction across all constructed FLS networks in OA and RA) to its TGs into 4 separate quartiles (Q1-Q4, Supplementary Figure S2). We anticipated that the expression of TGs with a high regulatory weight (Q4) would be more significantly affected by a BACH1 knockdown than those with a low regulatory weight (Q1). To investigate this hypothesis, we used siRNA to knockdown BACH1 in RA FLS cell lines and performed RNA sequencing to assess the resulting changes in gene expressions (Methods 2.8). We identified 24 up-regulated and 14 down-regulated genes between the two groups (Student’s *t* test, *p <* 0.05 after Benjamini-Hochberg correction [61], Figure 5A, Supplementary Table S13). Interestingly, BACH1’s target genes with the lowest edge weight (Q1) had a significantly lower measured fold-change compared to those with the highest weights (Q4) (0.3 vs 0.5, *p* = 3.46 *×* 10*^−^*^7^, Student’s t-test), indicating a good agreement between the measured gene expression fold-changes with the regulatory edge weights of the BACH1 FLS networks (Figure 5B), further validating the RA FLS network we constructed.

**Figure 5:**
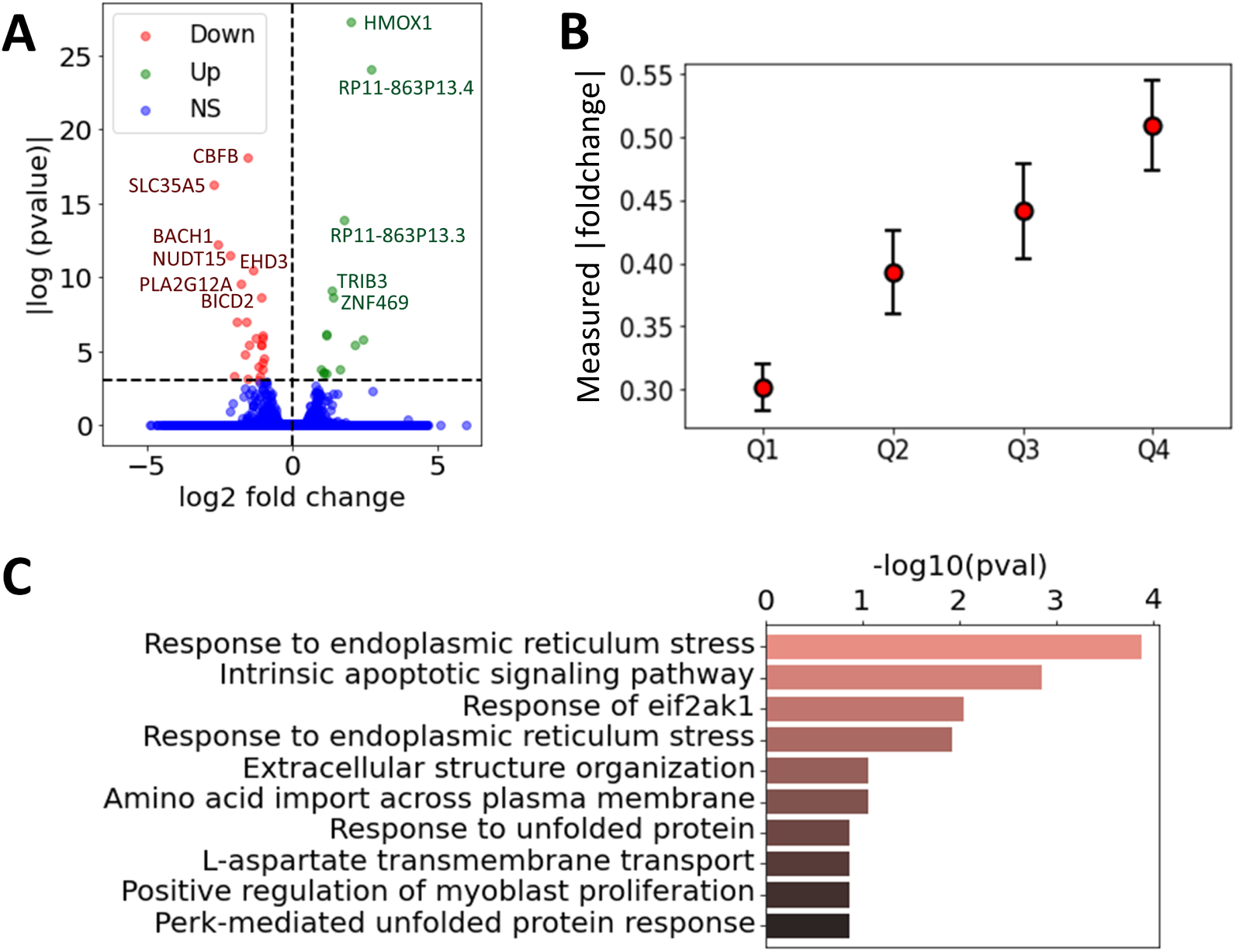
(A) Volcano plot of the measured log2 fold change and *p*-values for all the expressed genes without and with BACH1 knockdown (paired Student’s *t*-test), where the genes with a *p*-values below 0.001 were annotated. Genes were labeled as downregulated, upregulated, or non-significant (NS). (B) Measured *|*log2 foldchange*|* of gene expression between the siBACH1 and siCTL group, averaged over BACH1 target <u>gene</u>s grouped by the four quartiles Q1-Q4. Errors bars represent the 95% confidence interval, defined as *std*/ *|Q_n_|*. (C) Top 10 over-represented pathways identified among the differentialy expressed genes with and without BACH1 knockdown.

Among the genes with the most significant increase in mRNA levels after siRNA BACH1 knockdown were HMOX1, RP11-863P13.3 and RP11-863P13.4 (two lincRNAs), ZNF469 and TRIB3, and among those with reduced expression was SLC35A5 (a nucleoside-sugar transporter), NUDT15, CBFB (a TF) and STRADB (involved in cell polarity and energy-generating metabolism). Pathway analyses of the identified DEGs increased the representation of genes implicated in “intrinsic apoptotic signaling pathways in response to endoplasmic reticulum (ER) stress”, “response to ER stress” and “response to EIF2AK1” (Figure 5C, and Supplementary Table S14). BACH1 is implicated in oxidative stress [62], and oxidative stress may have a role in ER stress.

### 1.7 Effect of BACH1 knockdown on FLS migration, adhesion, lamellipodia formation, and cell morphology

We next examined the role of BACH1 in RA FLS behaviors relevant to disease and joint damage (Methods 2.9). siRNA knockdown of BACH1 significantly reduced RA FLS adhesion by an average of 50% (*p* = 0.021), and reduced FLS migration in the wound healing (scratch) assay by approximately 40% (*p* = 0.0067; paired *t*-test, Figure 6A). FLS cell morphology was analyzed under immunofluorescence microscopy, showcasing actin fibers and lamellipodia, marked by Phalloidin and pFAK staining, respectively (Methods 2.10, Figures 6B & C). Images were obtained from twenty cells per FLS cell line (a total of four different cell lines, each from a different RA patient). Knockdown of BACH1 significantly affected FLS morphology, reducing the number of cells with thick and polarized actin fibers, as well as reducing the number of cells with an elongated shape (Figure 6A & B). Knockdown of BACH1 also significantly reduced the numbers of pFAK positive lamellipodia, which has a critical role in movement and adhesion [63]. The FLS production of total or activated matrix metalloproteases (MMP2 and MMP3) and cell invasion through Matrigel were not significantly affected by BACH1 knockdown (Methods 2.11, Figure 6A and Supplementary Figure S3).

**Figure 6:**
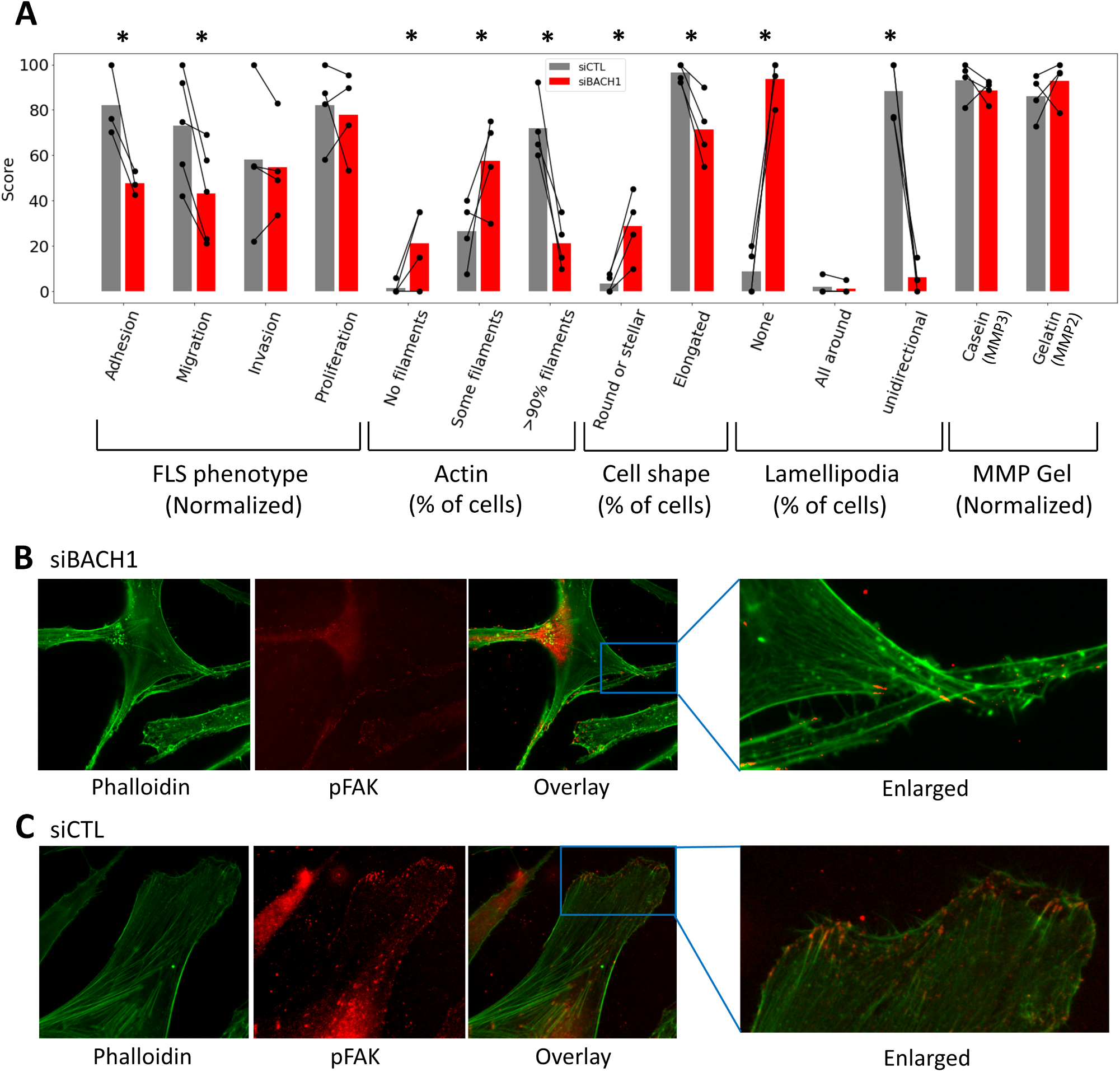
BACH1 silencing and changes in RA FLS. (A) BACH1 silencing reduced RA FLS adhesion, migration (wound healing assay), and affected cell morphology, including key characteristics required for movement and invasion such as thick and linear actin filaments, elongated shape, and the unidirectional formation of lamellipodia (bars represent the mean of each phenotype in each group; * *p <* 0.05; paired *t*-test). Phenotype scores on the *y*-axis were quantified either by the % of cells with the described attribute, or by dividing the score by the highest measured value and multiplying by 100 (indicated as normalized). (B&C) Representative immunofluorescence microscopy images of RA FLS (magnification 500x), showing actin fibers and lamellipodia, marked by Phalloidin and pFAK, respectively. FLS were treated with (B) siRNA BACH1, showing a stellate morphology, with disorganized actin fibers and no lamellipodia, while (C) cells treated with siRNA control (CTL) had the typical RA FLS elongated morphology with thick and organized actin fibers as well as polarized formation of lamellipodia.

## Discussion

Sustained disease remission is still rarely achieved in RA [64]. Available treatments, including biologics and JAK inhibitors targeting different aspects of the immune response, achieve similar rates of response [65, 4]. FLS have a central role in RA pathogenesis [8], driving leukocyte chemotaxis into the synovial tissues, and mediating bone and cartilage damage [8]. The RA FLS have a highly invasive and destructive behavior that correlates with joint damage [66]. However, the FLS subtypes, states of activation, and their characteristics are only now beginning to be comprehended [11, 67, 20, 21], potentially paving the way for the identification of novel targets for the development of new treatments aiming to achieve sustained remission, while minimizing the risk of immunosuppression in RA patients. The recent availability of large transcriptomic studies and datasets have also expanded our understanding of RA pathogenic processes. Yet, the intricate molecular and cellular pathways, along with their regulation and interactions, remain largely elusive. Gene regulatory processes vary across cell types, and only a limited number of studies have characterized RA drivers in cell types relevant to RA pathogenesis, such as FLS [11, 68].

In this study, we leveraged FLS RNA gene expression data [20] to infer FLS-specific gene regulatory networks (GRNs), and created one for each synovial biopsy in the datasets. Then, a differential analysis of these networks enabled a comparison between RA vs OA and revealed key pathways and gene interactions associated with each condition. This analysis highlighted potential therapeutic targets and molecular markers that differentiate RA from OA. For example, we identified an overrepresentation of GO pathways involved in the regulation of GTPase activity in RA. Interestingly, GTPases such as Rac1 and RhoA have been implicated in the regulation of RA FLS behaviors including adhesion, migration, and invasion [69, 70, 71]. As these networks contained regulatory information between TFs and their TGs, our analysis allowed us to rank the TFs according to their contribution to the observed differential gene expression between RA and OA (Table 1).

We further refined our findings using an alternative computational methodology [36] to enhance robustness and reduce the number of false positives. With this approach, we identified 28 candidates key driver TFs in FLS (Table 2), including seven not previously implicated in RA (BACH1, HLX, ETV7, TGIF1, ELF1, HIVEP1, and PLAGL1). Of those seven, only BACH1 and HLX were specific to FLS, while the others were also associated with T and/or B cell signatures (Table 2). Interestingly, we identified significant TF-TF co-regulated networks where BACH1 was the most significant driver. Among the genes differentially targeted by BACH1 were genes involved in fatty acid degradation and ferroptosis, an iron-dependent programmed cell death. BACH1 promotes ferroptosis, in part by repressing genes that interfere with iron-induced oxidative stress [59, 72]. Ferroptosis has been implicated in cancer cell behavior and metastasis, but its potential role in RA remains uncharacterized. BACH1-induced ferroptosis has been recently demonstrated to inhibit fatty acid biosynthesis [60, 59], and fatty acid, essential for normal FLS functioning, are not as crucial for RA FLS, which have reduced usage and reduced fatty acid beta-oxidation[73]. BACH1 also regulates cancer cell metabolism beyond fatty acids, including the production of lactate via hexokinase 2 (HK2) [74]. HK2 is central to RA FLS behaviors and rodent arthritis [75]. Thus, BACH1 emerges as a novel regulator of both ferroptosis and metabolism in RA FLS.

Knock down of BACH1 in RA FLS significantly increased mRNA levels of HMOX1, a gene transcriptionally suppressed by BACH1. Other genes expressed in increased levels in BACH1 knockdown FLS included lincRNAs, the CBFB, and STRADB (involved in cell polarity and energy-generating metabolism). Knockdown of BACH1 increased the expression of “intrinsic apoptotic signaling pathways in response to endoplasmic reticulum (ER) stress” and “response to ER stress” pathways. Moreover, cell ER stress has been linked to driving synovial inflammation [76]. We examined the effect of siRNA knockdown on RA FLS phenotypes and observed that in the absence of BACH1, RA FLS were not able to take an elongated shape, and did not form thick actin fibers or lamellipodia. Without these morphological changes, FLS are unable to move, which is crucial for invasion. Indeed, BACH1 knockdown cells had a reduced ability to adhere and reduced mobility. Furthermore, BACH1’s role in osteoclastogenesis [77], combined with the critical function of osteoclasts in RA-induced bone damage, underscores the potential advantage of BACH1 inhibition in RA therapy.

In addition to the BACH1 discovery, the highest ranked TFs in RA FLS included two homeobox genes (NKX2-1 and HOXA1, Table 1) not previously implicated in arthritis. Our integrated strategy identified additional key TFs driving the gene expression signatures in FLS, and potentially new targets for functional studies including HLX, MITF, FOSL1, and ETV4, among others. FOSL1 and ETV7 were also among the top FLS specific TFs in a recent systems biology study incorporating epigenomic analyses [11], further validating our observations.

In conclusion, this work presents a novel and integrated methodology for analyzing RA FLS transcriptomic data. Our methodology led to the identification of new genes and pathways for further investigation and potential therapeutic targeting. We identified BACH1 as a new key TF driving the gene expression and phenotypic characteristics of RA FLS. Although BACH1 emerged as the primary driver in our analyses, our findings pave the way for additional investigations into the roles of various TFs in RA FLS and their co-regulation, and whether targeting one of them is enough to affect disease as BACH1 analyses suggest. In a broader context, the computational approach described in this article, with the use of statistical techniques to compare network properties across samples and phenotypic groups, has the potential to also effectively be utilized to analyze other synovial cell types, such as T cells, B cells, and monocytes [36], or be adapted to the study of other diseases.

## 2 Methods

### 2.1 Gene expression data and normalization

Our analysis leverages the cell-type specific bulk RNA-seq study of FLS from patients with RA (n = 18) and OA controls (n = 13) from the Accelerating Medicines Partnership (AMP) Phase I [20]. We did not use the scRNA-Seq data from this study as they only provided FLS scRNA-Seq expression from three patients. All data underwent scaling normalization [78] to remove potential biases of other experimental artifacts across samples. The assumption is that any sample-specific bias (e.g., in capture or amplification efficiency) affects all genes equally via scaling of the expected mean count for that gene. The size factor for each sample then represents the estimate of the relative bias in that sample, so division of its counts by its size factor should remove that bias.

### 2.2 Correlation of gene expression across cell types

For a given gene *g ∈ G*, after performing a Student’s *t*-test of its expression between RA and the control group in a given cell type *C*, we obtain its t-statistics denoted *t*_diff-expr_(*g*, *C*). Then we compute the correlation of this score across a cell type pair (*C*1, *C*2) as:

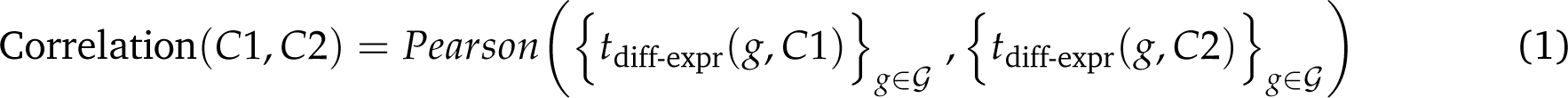

### 2.3 Gene regulatory networks in FLS

We inferred our gene regulatory networks with PANDA [79] by combining gene expression profiles of FLS [20] with priors knowledge about TF binding motifs (binary) and TF-TF interactions [80, 81]. These were inferred from the StringDB [32] and CIS-BP database [31], and can be downloaded directly from the GRAND database [82] (https://grand.networkmedicine.org/). Briefly, PANDA uses message passing to integrate a prior network (obtained by mapping TF motifs to the genome) with protein-protein interaction and gene expression data by optimizing the weights of edges in the networks with iterative steps. Applied to our data, PANDA produced directed networks of TFs to their TGs, comprising 644 TFs and 18 992 genes, resulting in 12 230 848 edges. Here, each edge between a TF and its TG is associated a weight, which represent the likelihood of a regulatory interaction between the TF and its TG. The weight values are normalized with a Zscore and range roughly from -3 and 3, which corresponds to how many standard deviation it is below (negative Zscore) or above (positive Zscore) the mean of all other weight in the network.

Then, we used LIONESS [33] to estimate an individual gene regulatory network for each sample in the population. LIONESS estimates sample specific networks by sequentially leaving each sample out, calculating a network (with PANDA) with and without that sample, and using linear interpolation to estimate the network for the left-out sample. All networks were inferred with python library netZooPy (https://github.com/netZoo/netZooPy).

### 2.4 Analysis of TFs RA regulatory activity in gene regulatory networks

For each individual sample in our RNA-seq data, we established a FLS-directed network of TFs nodes with regulatory edges linking them to their TGs, with weight representing the likelihood of the regulatory interaction between the two nodes. We leveraged this collection of networks to test whether the weights of these regulatory edges differed significantly between RA and control tissues, and to identify the TFs driving these regulatory differences. The Student’s t-test was used for (i) the differential gene expression between RA and control group and (ii) the differential weight of the regulatory edges between RA and control group. The obtained scores are denoted as *t*_diff-expr_ and *t*_diff-edge_, respectively. We define RA DEGs the genes having a *|t*_diff-expr_*| >* 1 in their RA differential expression. Note that from the definition of the *t*-score, these represent genes where the difference between the two phenotype groups is higher than the standard deviation of their gene expression across all samples. Then, we quantified the TFs regulatory importance as the average absolute differential weight of the regulatory edges *|t*_diff-edge_*|* of the RA targeted genes, where only the TGs listed in the TF motifs were considered (Eq. 2). We expected that TFs with the highest scores would be the most likely to contribute to RA regulation. Defining *G* as the set of all genes in the network, we formalize the computation of the TF scores with

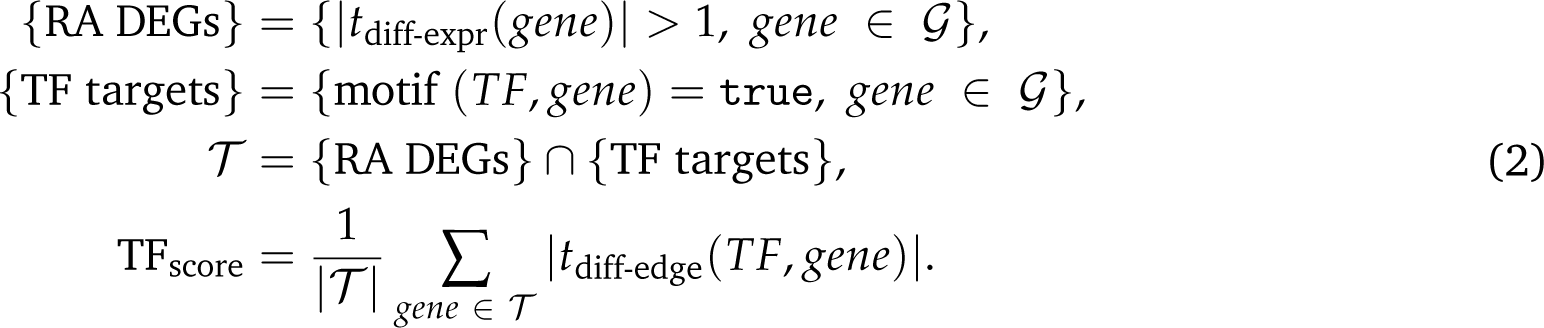

### 2.5 TF-TF co-regulation network

We quantify the co-regulation between TFs by evaluating the Pearson correlation between their common gene target’s differential edge weight. TFs with less than 10 common target are associated to co-regulation of 0. Defining *G* the set of all genes in the network, we write

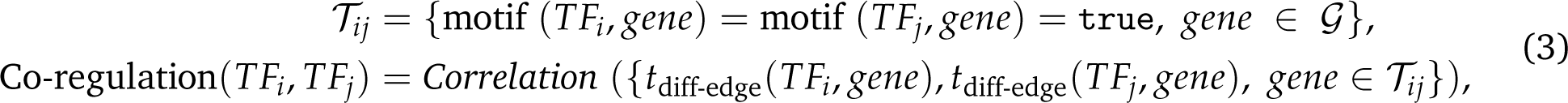

### 2.6 Key driver analysis

Two independent list of RA associated genes, denoted as the *DEG list* and *Literature list*, were compiled with a DEG meta analysis [83, 84, 85, 86, 87, 87] and by aggregating together several databases [88, 89, 90, 16, 91, 92, 93, 94], respectively (Supplementary Table S7). Then, 14 networks from different human organs, tissues and cell types were downloaded (Supplementary Table S8). Each of these networks and list of RA associated genes were used to run a key driver analysis (KDA) with the mergeomics R library [35]. For additional details, refer to [36].

### 2.7 Isolation and culture of FLS

RA FLS were obtained as previously described [71, 95]. Briefly, synovial tissues were obtained under IRB-approved protocols and all patients signed informed consent forms. Tissues were freshly obtained, minced and incubated with a solution containing DNase (0.15 mg/mL), hyaluronidase type I-S (0.15 mg/mL) and collagenase type IA (1 mg/mL) (Sigma) in DMEM (Invitrogen, Carlsbad, CA, USA) for 1 hour at 37°C. Cells were washed and re-suspended in complete media containing DMEM supplemented with 10% FBS (Invitrogen), glutamine (300 ng/mL), amphotericin B (250 µg/mL) (Sigma) and gentamicin (20 µg/mL) (Invitrogen). After overnight culture, non-adherent cells were removed and adherent cells cultured. All experiments were performed with FLS after passage four (>95% FLS purity).

### 2.8 siRNA knockdown

RA FLS (4-5 cell lines from different patients) were transfected with siRNA BACH1 or a non-coding control using the Dharmacon SMARTpool siRNA according to the manufacturer’s instructions (Dharmacon, GE Lifesciences, Lafayette, CO, USA) as previously described [71]. Cells were then incubated at 37°C for 24-48 hours prior to initiating the functional assays described bellow, or using the cells for RNA sequencing. Knockdown was confirmed with qPCR.

### 2.9 FLS assays

#### Invasion assay

The in vitro invasiveness of FLS was assayed in a transwell system using Matrigel-coated inserts (BD Biosciences, Franklin Lakes, NJ, USA), as previously described [67, 95, 14]. Briefly, siRNA transfected FLS were harvested by trypsin-EDTA digestion and re-suspended in 500 *µ*L of serum-free DMEM. 2×104 cells were placed in the upper compartment of each Matrigel-coated inserts. The lower compartment was filled with media containing 10% FBS and the plates incubated at 37°C. After 24 hours, the upper surface of the insert was wiped with a cotton-swab to remove non-invading cells and the Matrigel layer. The opposite side of the insert was stained with Crystal Violet (Sigma) and the number of cells that invaded through Matrigel analyzed with Image J software. Experiments were done in duplicate.

#### Adhesion to Matrigel assay

The FLS adhesion assay was done as previously reported [70]. Briefly, transfected cells were trypsinized and counted. Six thousand cells per well were plated in triplicate in 96-well plate previously coated with 5 *µ*g/mL of Matrigel (BD), in complete media. After two hours non-adherent cells were wash out with PBS 1X, and adherent cells stained with Crystal Violet. Cell were manually counted and read with spectrophotometer at 590 nm.

#### Migration in the wound healing assay

Transfected FLS were briefly trypsinized and counted. Six thousand cells per well were plated in triplicates in a 96-well plate. Cells were allowed to grow untill confluent (usually 24 hours). Then, a wound (scratch) was created by using a 10 *µ*L pipette tip. Pictures were taken at this initial time (time 0) and 24 hours later. FLS migration was determined using image J software by subtracting the density (the number of cells that cross into the wound/scratch area) after 24 hours from the density of time of time 0 (reference point).

#### Proliferation

Transfected FLS were trypsinized and counted. Three thousand cells per well were plated in triplicates in a 96-well plate in complete media with 10% FBS. After the indicated times, cells were incubated with Promega^™^ CellTiter 96^™^ AQueous One Solution Cell Proliferation Assay (MTS) (Madison, WI) according to the manufacturer’s instructions. Proliferation was assessed by colorimetric reading at 490 nm.

### 2.10 Immunofluorescence microscopy

Immunofluorescence was performed as previously reported [70]. Briefly, siRNA transfected cells were plated on glass coverslip with media containing 10% FBS. Cells were fixed with 4% formaldehyde for 15 min at room temperature and permeabilized with PBS/Triton X-100 0.1% for 5 min. Non-specific binding was blocked with 5% nonfat milk. Cells were then stained with Alexa Fluor 488 (green) Phalloidin (Invitrogen) to stain the actin filament, and anti-phospho-FAK (pFAK; Abcam, Cambridge, Massachusetts, USA), followed by a secondary Alexa Fluor 594 (red) antibody to identify pFAK and lamellipodia. Images were acquired with a Leica DMi8 microscope at 600× magnification and analyzed with the Leica application suite X (LAS X) software (Leica).

### 2.11 Matrix metalloprotease (MMP) quantification in zymograms

Gelatin (MMP2) and casein (MMP3) zymography was performed according to previously described methods [14]. Briefly, RA FLS transfected with either siRNA BACH1 or control were cultured on Matrigel and supernatants concentrated with Micron centrifugal filters (Millipore Sigma) and the protein content quantified. The same amount of protein per samples was used in each experiment. Protein was mixed with Tris–glycine–sodium dodecyl sulfate (SDS) sample buffer (Invitrogen), loaded into a zymogram pre-casted gel (Invitrogen), and run for 90 minutes at 125V. After electrophoresis, gels were treated with renaturing buffer (Invitrogen), followed by incubation in developing buffer (Invitrogen) at 37*^◦^*C overnight. Gels were stained with SimplyBlue SafeStain (Invitrogen) for 1 hour at room temperature and washed. Areas of protease activity appeared as clear bands against a dark-blue background.

### 2.12 RNA extraction and RNA sequencing

Total RNA were extracted and isolated from synovial tissues, quantified by Nanodrop and 400 ng per sample and sent to Novogene (Beijing, China) for sequencing and analyses.

## Data availability

All the gene lists obtained in this study, along with the data and the code to reproduce all figures presented in this article are made available publicly on Github at https://github.com/AI-SysBio/ RA-drug-discovery. PANDA and LIONESS networks generated in this study will be uploaded upon acceptance.

## Acknowledgments and Funding

This research was supported by the COSMIC European Training Network, funded from the European Union’s Horizon 2020 research and innovation program under grant agreement No 765158. Dr. Gulko and Laragione were funded by the NIH R01AR07316, by the Icahn School of Medicine at Mount Sinai, and by the Eunice Bernhard fund.

## Author contributions statement

A.P. performed all computational analysis, under the supervision of M.R.M and P.G. T.L. performed the experiments related to the silencing of BACH1, under the supervision of P.G. All authors participated in the regular discussions and reviewed the manuscript.

## Supplementary Materials

### A Supplementary Tables

**Table S1:** Top 30 differentially expressed genes in FLS RA *vs* OA, ranked by *|t*-score*|*. The *p*-values were not adjusted.

| Gene (top 15) | $t$-score | $p$-value | Gene (rank 16-30) | $t$-score | $p$-value |
| --- | --- | --- | --- | --- | --- |
| LINC01600 | 4.70 | $6.22 \times 10^{-5}$ | CPAMD8 | -3.47 | $1.70 \times 10^{-3}$ |
| SLC25A26 | 4.29 | $1.94 \times 10^{-4}$ | PRKG1 | 3.45 | $1.79 \times 10^{-3}$ |
| MMD | -3.90 | $5.46 \times 10^{-4}$ | RAB15 | -3.38 | $2.14 \times 10^{-3}$ |
| NUDT21 | -3.77 | $7.74 \times 10^{-4}$ | ACYP2 | 3.37 | $2.20 \times 10^{-3}$ |
| ATP8A1 | -3.76 | $7.94 \times 10^{-4}$ | TRDN | 3.37 | $2.23 \times 10^{-3}$ |
| GPR4 | 3.74 | $8.49 \times 10^{-4}$ | SOAT2 | 3.37 | $2.23 \times 10^{-3}$ |
| ARHGAP28 | 3.68 | $9.84 \times 10^{-4}$ | NFATC3 | 3.36 | $2.29 \times 10^{-3}$ |
| BIRC7 | 3.67 | $1.01 \times 10^{-3}$ | E2F5 | 3.34 | $2.39 \times 10^{-3}$ |
| ALOX15B | -3.63 | $1.13 \times 10^{-3}$ | EGFL7 | -3.33 | $2.47 \times 10^{-3}$ |
| STKLD1 | -3.58 | $1.28 \times 10^{-3}$ | MUSTN1 | 3.32 | $2.53 \times 10^{-3}$ |
| TMEM128 | 3.58 | $1.29 \times 10^{-3}$ | TRHDE | 3.32 | $2.54 \times 10^{-3}$ |
| E2F1 | 3.55 | $1.38 \times 10^{-3}$ | FAM47E | 3.31 | $2.56 \times 10^{-3}$ |
| PRKDC | -3.53 | $1.47 \times 10^{-3}$ | ANKRD11 | -3.30 | $2.61 \times 10^{-3}$ |
| PROS1 | 3.50 | $1.58 \times 10^{-3}$ | TMEM154 | -3.30 | $2.62 \times 10^{-3}$ |
| CNFN | -3.47 | $1.69 \times 10^{-3}$ | SAMD4A | -3.30 | $2.62 \times 10^{-3}$ |

**Table S2:** Top10 Pathway of FLS DEGs (1093 genes).

| Biological Process/Pathway | Genes | adj. *p*-value |
| --- | --- | --- |
| Negative regulation of low-density lipoprotein receptor activity. | ABCA2, APP, ADIPOQ, PCSK9, ITGAV (5/7) | 0.035 |
| Arrhythmogenic right ventricular cardiomyopathy. | SLC8A3, DES, CDH2, SGCA, ITGA3, ITGA11, ITGB8, CACNA2D2, ITGAV, ITGA6, CACNA1D, ITGA5 (12/77) | 0.231 |
| Ubiquinone and other terpenoid-quinone biosynthesis. | NQO1, COQ2, TAT, COQ6 (4/11) | 0.231 |
| Axon guidance. | NTNG2, ROBO3, SEMA6D, SEMA3A, SEMA3B, NFATC3, PIK3CD, UNC5C, SEMA3E, SSH3, RHOD, RGMA, EFNB2, EFNA3, EFNB3, ABLIM3, RAC2, EPHB2, MET, PLXNA4 (20/182) | 0.231 |
| Hypertrophic cardiomyopathy. | SLC8A3, PRKAA2, DES, SGCA, ITGA3, ITGA11, ITGB8, CACNA2D2, ITGAV, ITGA6, CACNA1D, ITGA5 (12/90) | 0.265 |
| Arginine and proline metabolism. | GAMT, MAOB, CKM, SMOX, PYCR2, HOGA1, PRODH, L3HYPDH (8/50) | 0.299 |
| Dilated cardiomyopathy. | SLC8A3, PLN, DES, SGCA, ITGA3, ITGA11, ITGB8, CACNA2D2, ITGAV, ITGA6, CACNA1D, ITGA5 (12/96) | 0.299 |
| Carnitine shuttle. | PRKAA2, CPT2, THRSP, ACACB, ACACA (5/11) | 0.317 |
| Regulation of vascular associated smooth muscle cell migration. | FGF9, IGFBP5, ADIPOQ, DOCK7, PRKG1 (5/12) | 0.345 |
| ECM-receptor interaction. | LAMA5, SV2B, TNN, ITGA3, ITGA11, ITGB8, ITGAV, ITGA6, ITGA5, COL9A2, THBS1 (11/88) | 0.353 |

**Table S3:** List of the top TF regulators in FLS with a Z-statistics above 1, provided in parenthesis.

| Rank 1-25 | Rank 26-50 | Rank 51-75 | Rank 76-91 |
| --- | --- | --- | --- |
| RORC (3.39) | MNT (1.87) | ZBTB4 (1.49) | ATF2 (1.20) |
| NKX2-1 (3.37) | BACH1 (1.83) | HNF1B (1.48) | GSC2 (1.20) |
| HOXA1 (3.20) | IKZF1 (1.82) | IRX1 (1.47) | NFYB (1.18) |
| ETV2 (2.67) | FEZF1 (1.82) | DPRX (1.45) | IRF7 (1.18) |
| MITF (2.64) | SOX10 (1.81) | HLF (1.45) | CREM (1.16) |
| SIX5 (2.61) | PGR (1.76) | IRF6 (1.43) | MEIS2 (1.16) |
| ELF5 (2.46) | CREB3 (1.75) | TFCP2 (1.43) | NFAT5 (1.15) |
| ZBTB1 (2.42) | FOS (1.72) | TGIF2LX (1.42) | SOX30 (1.15) |
| FOXC2 (2.27) | CEBPZ (1.66) | HOXB8 (1.42) | GATA2 (1.14) |
| TBX4 (2.23) | HOXB6 (1.66) | GLIS1 (1.39) | NFATC1 (1.13) |
| CBFB (2.16) | MEOX2 (1.64) | ZNF713 (1.39) | FOSL1 (1.12) |
| BBX (2.16) | GABPA (1.64) | NFE2 (1.38) | CEBPB (1.11) |
| TAL1 (2.13) | MEIS1 (1.63) | THRB (1.34) | FOXO4 (1.09) |
| ZBTB3 (2.11) | HOXB1 (1.63) | ELF4 (1.34) | EN2 (1.09) |
| RARG (2.09) | NR6A1 (1.62) | HLTF (1.32) | FLI1 (1.09) |
| ETV3 (2.04) | E4F1 (1.61) | TGIF1 (1.29) |  |
| TLX3 (1.97) | ELF3 (1.60) | HOXB5 (1.26) |  |
| HOXB3 (1.95) | CDX1 (1.58) | RUNX1 (1.27) |  |
| PKNOX1 (1.95) | ATF7 (1.58) | NPAS4 (1.26) |  |
| MECOM (1.94) | CLOCK (1.58) | BARX2 (1.25) |  |
| ELK4 (1.92) | ETV7 (1.57) | HOXA2 (1.25) |  |
| HOXB7 (1.91) | RFX5 (1.54) | E2F4 (1.25) |  |
| DBX2 (1.91) | HIF1A (1.52) | GCM2 (1.24) |  |
| HLX (1.90) | FOXI1 (1.50) | ZKSCAN3 (1.21) |  |
| TBX2 (1.90) | TBX5 (1.50) | VTN (1.21) |  |

**Table S4:** Top10 Pathway of FLS top TFs (185 TFs).

| Biological Process/Pathway | Genes | adj. p-value |
| --- | --- | --- |
| Activation Of HOX Genes During Differentiation. | RARG, MEIS1, HOXA2, HOXB3, HOXA1, HOXB1, PKNOX1 (7/91) | 0.000030 |
| Regulation of myeloid cell differentiation. | NFE2, MEIS1, CBFB, HOXB8, MEIS2, RUNX1 (6/68) | 0.000037 |
| Anterior/posterior pattern specification. | CDX1, HOXB3, HOXB7, HOXB6, HOXB5 (5/63) | 0.000513 |
| Developmental Biology. | CEBPB, RARG, CBFB, HNF1B, GATA2, SOX10, FLI1, RUNX1, MEIS1, NR6A1, TAL1, HOXA2, HOXB3, HOXA1, HOXB1, PKNOX1 (16/1073) | 0.000764 |
| Embryonic limb morphogenesis. | RARG, TBX5, TBX4, TBX2 (4/36) | 0.000942 |
| Regulation of antigen receptor-mediated signaling pathway. | ELF1, CBFB, RUNX1 (3/13) | 0.001054 |
| Human T-cell leukemia virus 1 infection. | FOSL1, ELK4, ATF2, CREB3, NFYB, NFATC1, FOS (7/219) | 0.003470 |
| Regulation of cardiac muscle cell proliferation. | TBX5, RUNX1, TBX2 (3/21) | 0.003834 |
| Regulation of response to cytokine stimulus. | ELF1, CBFB, RUNX1 (3/22) | 0.003834 |
| Negative regulation of myeloid cell differentiation. | MEIS1, HOXB8, MEIS2 (3/22) | 0.003834 |

**Table S5:** Top 100 TGs in FLS. The number in parenthesis corresponds to the number of key TF drivers (182 TFs with Z-statistics *>* 0.5) differentially targeting this gene (|*t*_diff-edge_|>1). Note that only genes differentially expressed between RA and OA FLS are considered here, |*t*_diff-expr_|>1.

| Rank 1-25 | Rank 26-50 | Rank 51-75 | Rank 76-100 |
| --- | --- | --- | --- |
| ALDH1A2 (67) | PAPOLA (44) | POT1 (39) | MET (34) |
| SYNE2 (64) | EFL1 (43) | LPP (39) | SORBS2 (34) |
| NF1 (62) | UBR4 (43) | ULK2 (38) | SHANK2 (33) |
| FOCAD (59) | USP34 (43) | RNF220 (38) | GALC (33) |
| EML1 (58) | DLG2 (43) | DPYS (38) | RBMS1 (33) |
| ABCA5 (56) | ZNF532 (42) | SLC9B2 (38) | RMDN1 (33) |
| DCDC1 (54) | TFDP2 (42) | PAH (38) | ANGPT1 (33) |
| TRPM8 (53) | CD163L1 (42) | EYA4 (37) | MAP4K5 (33) |
| COBLL1 (52) | TACC2 (42) | ITGAV (37) | CPNE4 (33) |
| HDAC6 (50) | MPHOSPH9 (42) | SFPQ (37) | CYLD (32) |
| VPS8 (50) | GTDC1 (44) | NUP205 (37) | CTNNA3 (32) |
| USP47 (49) | EFCAB5 (44) | ANKRD26 (67) | BCAS3 (32) |
| ASAP1 (49) | TM7SF3 (44) | HERC4 (36) | XPNPEP1 (32) |
| NOSTRIN (48) | ELAPOR2 (44) | RBM17 (36) | PHTF1 (32) |
| TCF4 (48) | ABI3BP (44) | UVRAG (35) | TRIM29 (32) |
| PTPN12 (47) | FRMD3 (44) | TNIK (35) | JKAMP (32) |
| MSI2 (47) | GRAMD2B (40) | MYBPC1 (35) | TNIP3 (31) |
| CLTC (46) | DNAH14 (40) | TAOK3 (35) | MAN2A1 (31) |
| DENND2B (46) | UTRN (40) | FN1 (34) | MAPK6 (31) |
| ADGRA3 (45) | SKIL (40) | SLC11A2 (34) | REXO2 (31) |
| SOX6 (45) | ROBO2 (40) | STPG1 (34) | SEMA6B (31) |
| CBWD5 (44) | NFASC (39) | RPRD1A (34) | NCAPD3 (31) |
| ANK3 (44) | SEMA6A (39) | ATP6V0A1 (34) | DDX46 (31) |
| SIPA1L1 (44) | EHBP1 (39) | GON4L (34) | TCERG1 (31) |
| TUT4 (44) | ATAD2B (39) | TLN2 (34) | CBWD6 (31) |

**Table S6:** Top10 Pathway of FLS top TGs (211 genes).

| Biological Process/Pathway | Genes | adj. p-value |
| --- | --- | --- |
| Positive regulation of GTPase activity. | BCR, BCAS3, SIPA1L1, DOCK9, TBC1D5, NF1, PIP5K1A, ITGA6, SIPA1L2, MAPRE2, RGS6, MAP4K4 (12/214) | 0.004845 |
| Cell-substrate junction assembly. | BCR, ACTN2, FN1, PIP5K1A, ITGA6 (5/34) | 0.017964 |
| Regulation of GTPase activity. | BCAS3, FGD5, SIPA1L1, DOCK9, RDX, NF1, ITGA6, MAPRE2, RGS6, MAP4K4 (10/189) | 0.017964 |
| Plasma membrane bounded cell projection morphogenesis. | TAOK3, DTNBP1, TNIK, MYO16, MAP4K4 (5/52) | 0.086533 |
| Negative regulation of chemotaxis. | ROBO2, SEMA6B, SEMA6A, PTPN2 (4/32) | 0.096032 |
| Regulation of endothelial cell migration. | BCAS3, ANGPT1, NF1, PTPRM, PRCP, FOXP1 (6/89) | 0.096032 |
| Focal adhesion assembly. | BCR, ACTN2, PIP5K1A (3/21) | 0.108547 |
| Nucleus organization. | SUN3, NUP205, TPR, LEMD3 (4/48) | 0.108547 |
| Positive regulation of focal adhesion disassembly. | MAPRE2, MAP4K4 (2/6) | 0.108547 |
| Retinal ganglion cell axon guidance. | ROBO2, PTPRM (2/6) | 0.108547 |

**Table S7:**
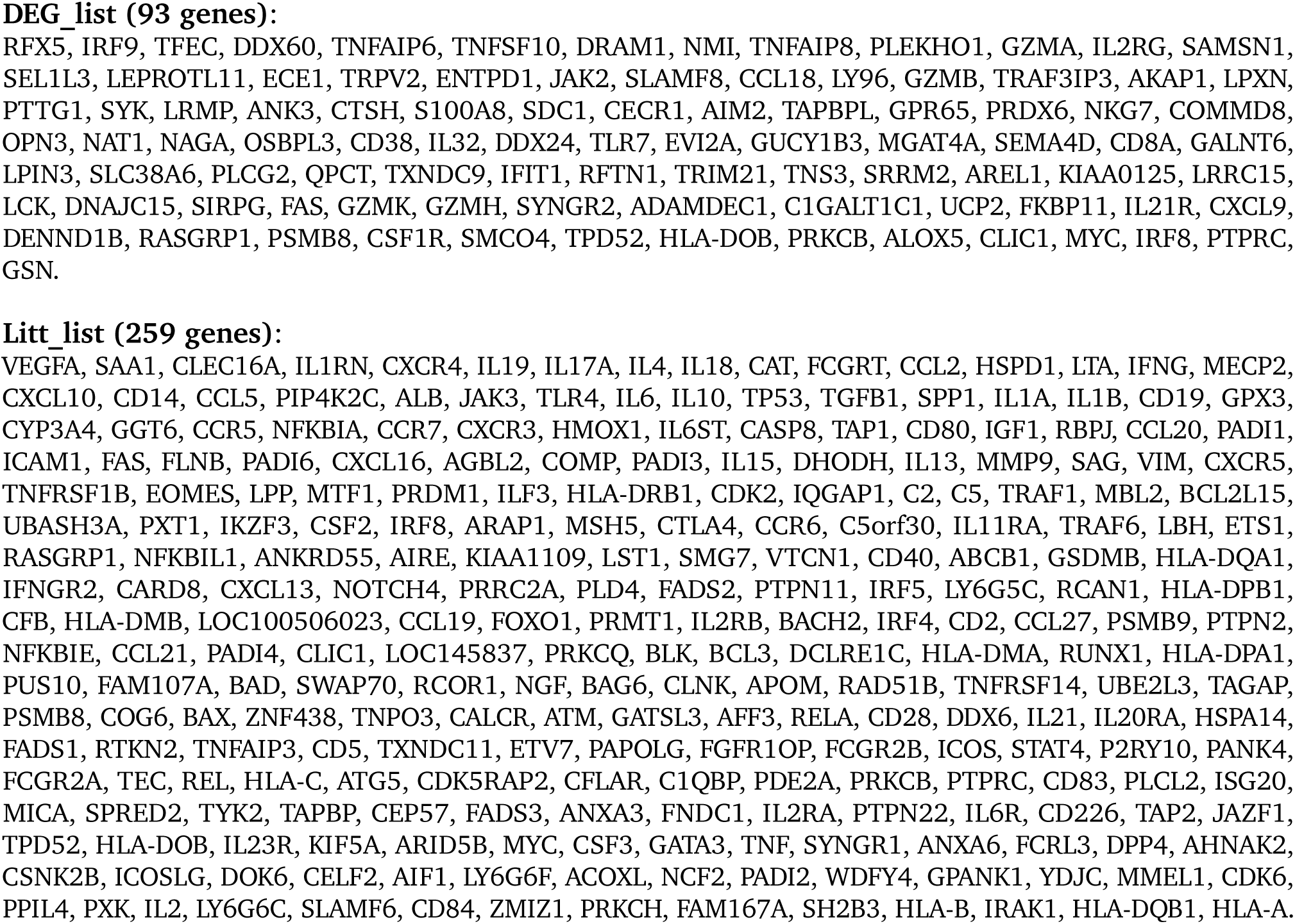
The two independent list of RA associated genes used in the Key Driver Analysis (KDA) [36].

**Table S8:** List of networks involved in our study. GIANT RIMBANET and PPI were downloaded from public database while PANDA and LIONESS networks were computed specifically for this study.

| Network(s) name | Type | Method | Used for | Number of networks |
| --- | --- | --- | --- | --- |
| PANDA_FLS | Bipartite | PANDA, LIONESS* | TF reg. score | 18RA, 12OA |
| GIANT_DC | Bayesian | GIANT<sup>§</sup> | KDA | 1 |
| GIANT_NKT | Bayesian | GIANT<sup>§</sup> | KDA | 1 |
| GIANT_Monocyte | Bayesian | GIANT<sup>§</sup> | KDA | 1 |
| GIANT_Tcell | Bayesian | GIANT<sup>§</sup> | KDA | 1 |
| GIANT_Bcell | Bayesian | GIANT<sup>§</sup> | KDA | 1 |
| GIANT_Fibroblast | Bayesian | GIANT<sup>§</sup> | KDA | 1 |
| GIANT_Adipocyte | Bayesian | GIANT<sup>§</sup> | KDA | 1 |
| GIANT_Tonsil | Bayesian | GIANT<sup>§</sup> | KDA | 1 |
| GIANT_LN | Bayesian | GIANT<sup>§</sup> | KDA | 1 |
| GIANT_Blood | Bayesian | GIANT<sup>§</sup> | KDA | 1 |
| GIANT_Spleen | Bayesian | GIANT<sup>§</sup> | KDA | 1 |
| RIMBANET_Multitissue | Bayesian | RIMBANET<sup>†</sup> | KDA | 1 |
| PPI | Undirected | StringDB<sup>†</sup> | KDA | 1 |

\* Computed in this study from bulk RNAseq [20].
§ GIANT networks downloaded from https://hb.flatironinstitute.org/download.
† RIMBANET and PPI downloaded from http://mergeomics.research.idre.ucla.edu/samplefiles.php.

**Table S9:**
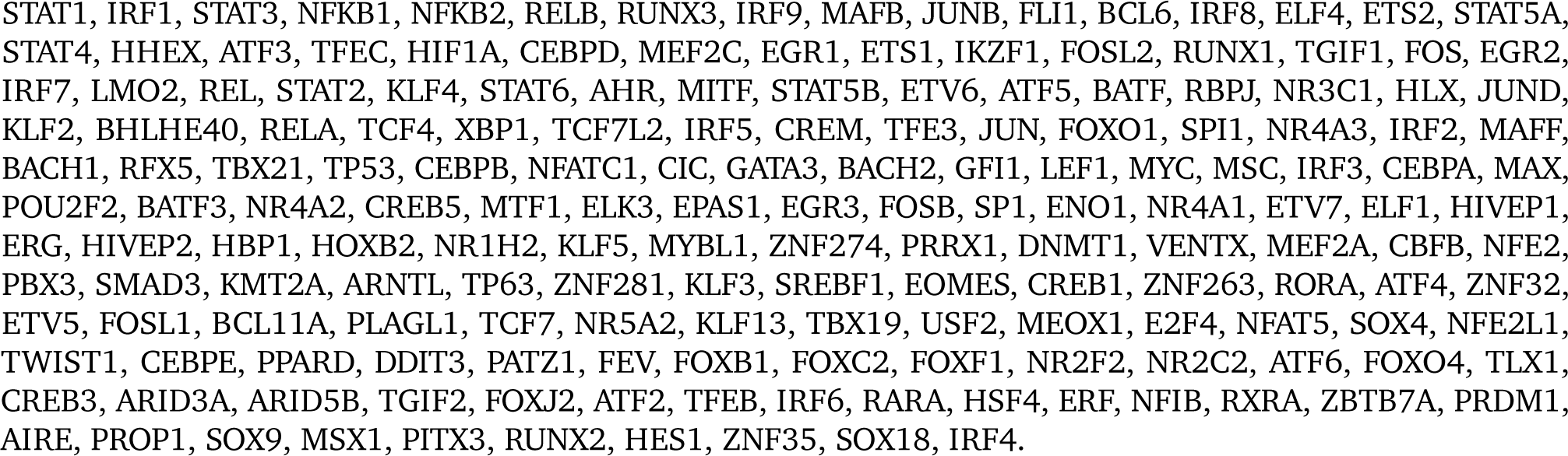
**list of KDA TF (174 TFs)** [36], ordered from highest to lowest score.

**Table S10:** The 28 key TFs implicated in the regulation of RA FLS, as identified in our analyses, are ranked by Z-statistics in each co-regulation cluster. The degree in the TF co-regulation network Figure 3A is also provided.

| TF | Cluster | FLS score<br>(dGRN Z-statistics) | Degree |
| --- | --- | --- | --- |
| BACH1 | 1 | 1.83 | 15 |
| FOS |  | 1.72 | 10 |
| HIF1A |  | 1.52 | 15 |
| TGIF1 |  | 1.39 | 17 |
| CREM |  | 1.16 | 13 |
| FOSL1 |  | 1.12 | 14 |
| FLI1 |  | 1.09 | 15 |
| BHLHE40 |  | 0.98 | 19 |
| STAT1 |  | 0.87 | 18 |
| EGR2 |  | 0.83 | 17 |
| NR4A1 |  | 0.75 | 17 |
| FOSL2 |  | 0.71 | 13 |
| JUNB |  | 0.68 | 14 |
| HIVEP1 |  | 0.65 | 16 |
| PLAGL1 |  | 0.62 | 21 |
| ENO1 |  | 0.59 | 15 |
| HLX | 2 | 1.90 | 12 |
| NFATC1 |  | 1.23 | 13 |
| IRF7 |  | 1.18 | 12 |
| HOXB2 |  | 0.54 | 3 |
| CBFB | 3 | 2.16 | 7 |
| RUNX1 |  | 1.27 | 8 |
| RFX5 |  | 1.54 | 2 |
| ETV7 | 4 | 1.57 | 10 |
| ELF1 |  | 1.03 | 8 |
| ELF4 |  | 1.34 | 4 |
| MITF | 5 | 2.64 | 10 |
| IKZF1 |  | 1.93 | 5 |

**Table S11:**
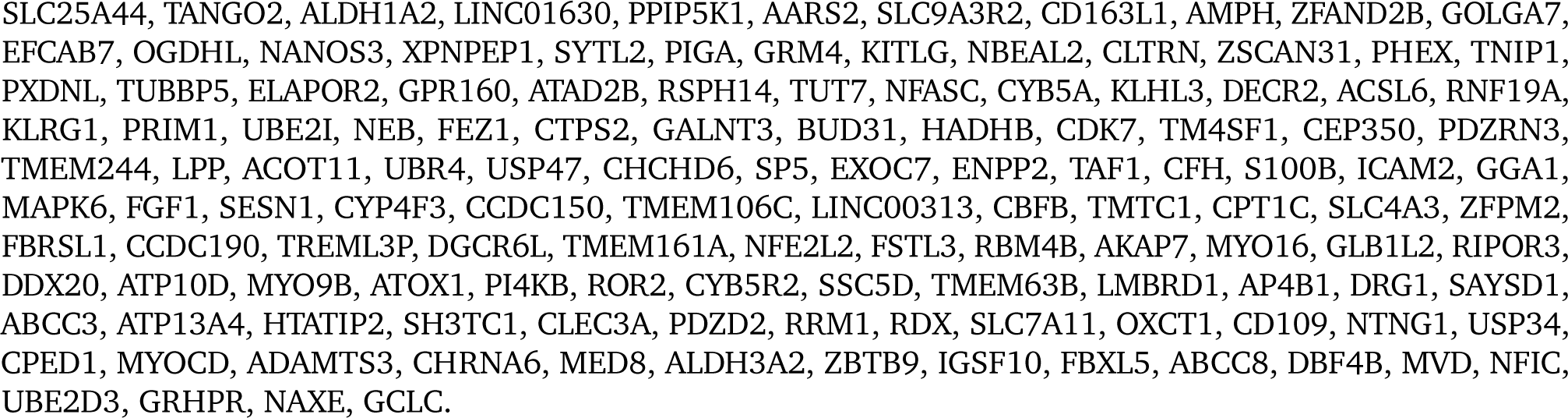
**List of BACH1 gene targets in the FLS network (131 genes)**, ranked by most to least differentially targeted in RA *vs* OA. Only genes with (|*t*_diff-edge_|>1 and |*t*_diff-expr_|>1) are shown in the list.

**Table S12:** Significant pathways of BACH1 targets (131 genes).

| Biological Process/Pathway | Genes | adj. *p*-value |
| --- | --- | --- |
| Fatty acid degradation. | ALDH3A2, HADHB, ACSL6, CPT1C (4/43) | 0.018154 |
| Ferroptosis. | GCLC, ACSL6, SLC7A11 (3/41) | 0.124481 |
| Valine, leucine and isoleucine degradation. | ALDH3A2, HADHB, OXCT1 (3/48) | 0.130240 |
| Metabolism of lipids. | ABCC3, UBE2I, CD163L1, CYP4F3, ACOT11, ACSL6, MED8, ALDH3A2, HADHB, OXCT1, MVD, PI4KB, DECR2 (13/732) | 0.261402 |
| Fatty acid metabolism. | ALDH3A2, HADHB, CYP4F3, ACOT11, ACSL6, DECR2 (6/173) | 0.261402 |
| Metabolism. | NAXE, CD163L1, PPIP5K1, OXCT1, ENPP2, DECR2, SLC25A44, ABCC3, CYB5A, RRM1, UBE2I, ABCC8, NANOS3, CYP4F3, ACOT11, ACSL6, MED8, CTPS2, ALDH3A2, HADHB, GRHPR, GCLC, LMBRD1, MVD, PI4KB (25/2049) | 0.269435 |

**Table S13:** Differentially expressed genes between siBACH1 and siCTL (adjusted *p*-value *>* 0.05).

| Gene (*downregulated*) | log2fold | *p*-value | Gene (*upregulated*) | log2fold | *p*-value |
| --- | --- | --- | --- | --- | --- |
| CBFB | -1.55 | $2.92 \times 10^{-12}$ | HMOX1 | 2.00 | $1.09 \times 10^{-16}$ |
| SLC35A5 | -2.71 | $2.44 \times 10^{-11}$ | RP11-863P13.4 | 2.72 | $5.19 \times 10^{-15}$ |
| BACH1 | -2.53 | $2.23 \times 10^{-9}$ | RP11-863P13.3 | 1.76 | $3.32 \times 10^{-10}$ |
| NUDT15 | -2.13 | $5.50 \times 10^{-9}$ | TRIB3 | 1.38 | $8.19 \times 10^{-8}$ |
| EHD3 | -1.33 | $1.65 \times 10^{-8}$ | ZNF469 | 1.40 | $1.57 \times 10^{-7}$ |
| PLA2G12A | -1.74 | $4.46 \times 10^{-8}$ | MIR22HG | 1.15 | $2.28 \times 10^{-6}$ |
| BICD2 | -1.07 | $1.47 \times 10^{-7}$ | SLC7A5 | 1.15 | $2.92 \times 10^{-6}$ |
| N6AMT1 | -1.55 | $9.09 \times 10^{-7}$ | RP11-863P13.5 | 2.45 | $4.40 \times 10^{-6}$ |
| STRADB | -1.90 | $9.16 \times 10^{-7}$ | TSPAN2 | 2.14 | $7.69 \times 10^{-6}$ |
| SYNM | -1.04 | $2.85 \times 10^{-6}$ | MEG9 | 1.63 | $5.07 \times 10^{-5}$ |
| FBXL17 | -1.25 | $3.54 \times 10^{-6}$ | SLC6A9 | 0.99 | $5.37 \times 10^{-5}$ |
| SLC2A3 | -1.00 | $3.96 \times 10^{-6}$ | CHAC1 | 1.09 | $6.35 \times 10^{-5}$ |
| C11orf95 | -1.05 | $6.61 \times 10^{-6}$ | KRTAP1-5 | 1.16 | $7.07 \times 10^{-5}$ |
| PRKAG2 | -1.08 | $7.00 \times 10^{-6}$ | ADM2 | 1.06 | $7.46 \times 10^{-5}$ |
| DICER1 | -1.48 | $7.65 \times 10^{-6}$ | | | |
| SENP1 | -1.62 | $1.56 \times 10^{-5}$ | | | |
| TNC | -0.99 | $2.14 \times 10^{-5}$ | | | |
| TOR1B | -1.03 | $2.89 \times 10^{-5}$ | | | |
| ARHGEF3 | -1.14 | $3.91 \times 10^{-5}$ | | | |
| RSPO2 | -1.02 | $4.70 \times 10^{-5}$ | | | |
| FAM134B | -1.10 | $9.65 \times 10^{-5}$ | | | |
| GOLT1B | -2.01 | $9.87 \times 10^{-5}$ | | | |
| MIB1 | -1.52 | $1.24 \times 10^{-4}$ | | | |
| DBP | -1.18 | $1.32 \times 10^{-4}$ | | | |

**Table S14:** Top10 Pathway analysis of siBACH1 silencing (740 genes).

| Biological Process/Pathway | Genes | adj. *p*-value |
| --- | --- | --- |
| Intrinsic apoptotic signaling pathway in response to endoplasmic reticulum stress. | PPP1R15A, DDIT3, ITPR1, TNFRSF10B, BAX, TRIB3, BAK1, CHAC1, BBC3, ATF4 (10/29) | 0.000134 |
| Intrinsic apoptotic signaling pathway. | PPP1R15A, ITPR1, IFI6, TNFRSF10B, TNFRSF1B, BBC3, HIPK2, DDIT3, BAX, TRIB3, CHAC1, BAK1, SCN2A, FNIP2, ATF4, EPHA2 (16/102) | 0.001431 |
| Response of EIF2AK1. | PPP1R15A, DDIT3, TRIB3, CHAC1, ATF5, ATF4 (6/15) | 0.009375 |
| Response to endoplasmic reticulum stress. | PPP1R15A, JUN, ITPR1, TNFRSF10B, THBS1, BBC3, HERPUD1, TMX1, DDIT3, BAX, TRIB3, CHAC1, BAK1, RNF185, ATF4 (15/110) | 0.012098 |
| Extracellular structure organization. | FBN2, COL15A1, ECM2, LAMC3, MMP1, ITGB3, TNC, F11R, THBS1, TGFBR1, COL1A1, ADAMTS15, COL3A1, SH3PXD2A, MMP16, COL5A3, SPP1, MMP19, COL9A3, ITGB6 (20/216) | 0.089010 |
| Amino acid import across plasma membrane. | SLC7A5, SLC6A9, SLC1A3, SLC1A4, SLC3A2, SLC1A5 (6/24) | 0.089010 |
| Response to unfolded protein. | HSPA8, HSPB8, DDIT3, HSPA4L, BAX, BAK1, TOR1B, HERPUD1 (8/49) | 0.139243 |
| L-aspartate transmembrane transport. | SLC1A3, SLC1A4, SLC1A5 (3/5) | 0.139243 |
| Positive regulation of myoblast proliferation. | ATF2, SOX15, GATA6 (3/5) | 0.139243 |
| PERK-mediated unfolded protein response. | IGFBP1, DDIT3, ATF4, HERPUD1 (4/11) | 0.139243 |

### B Supplementary Figures

**Figure S1:**
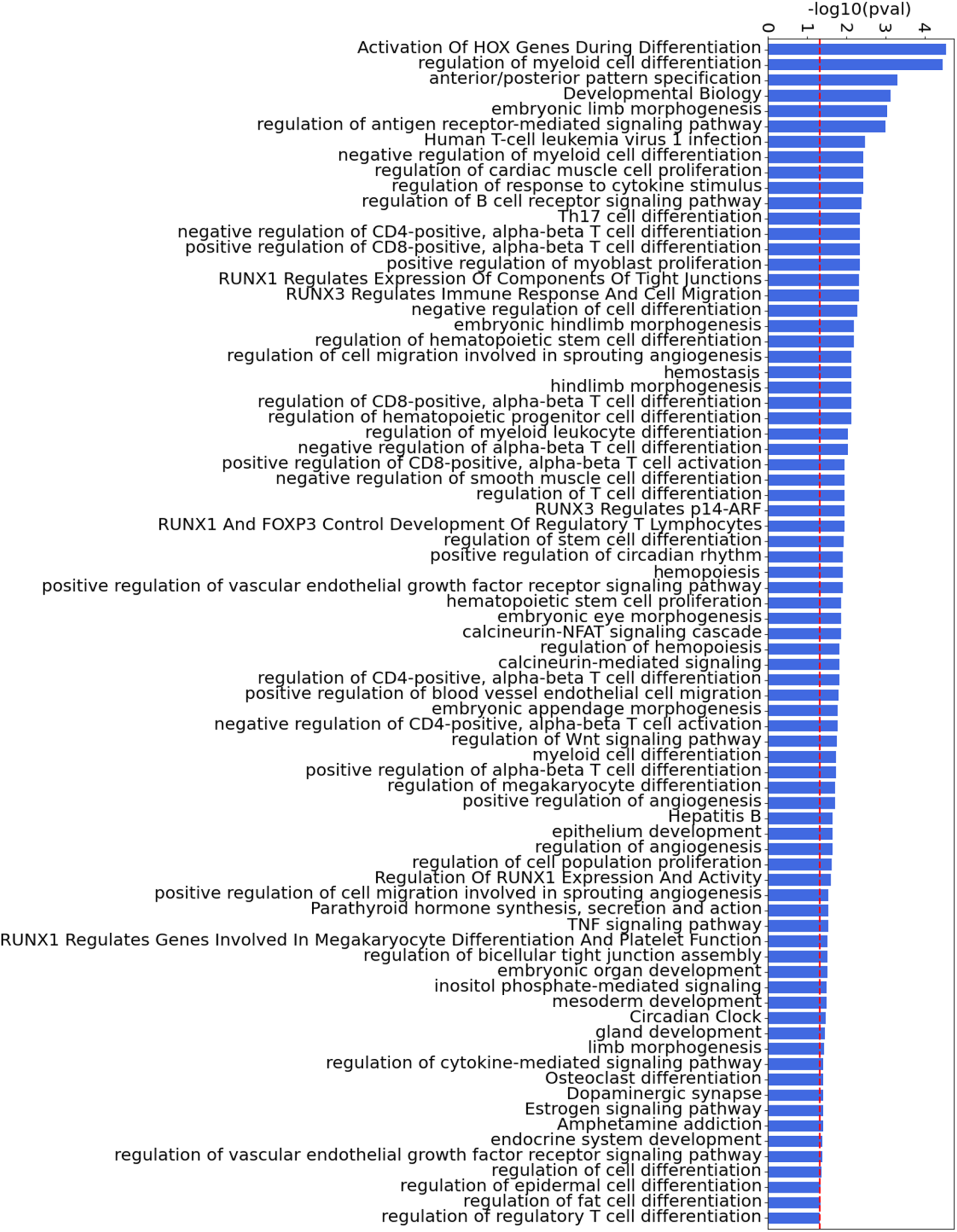
All significant pathways of the major TFs involved in FLS compiled from GO [28], KEGG [29] and Reactome [30], ranked by adjusted p-values. The threshold for significance (0.05) is shown as a dashed red line for visualization.

**Figure S2:**
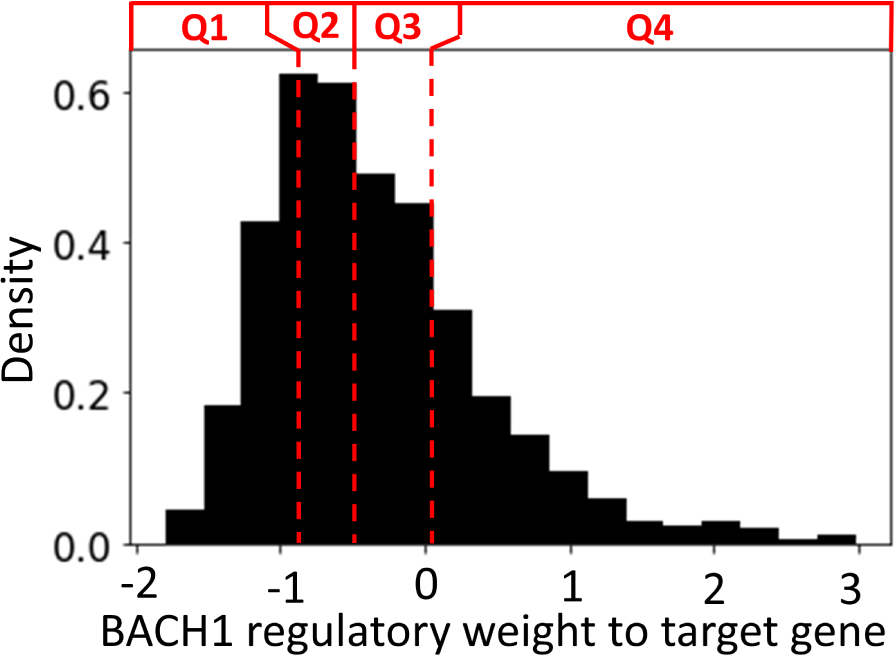
Density distribution of the edge weight values of BACH1 to its TGs, separated into four quartiles. Here, edge weight indicate the likelihood of interaction between BACH1 to its target, and is normalized with a Zscore.

**Figure S3:**
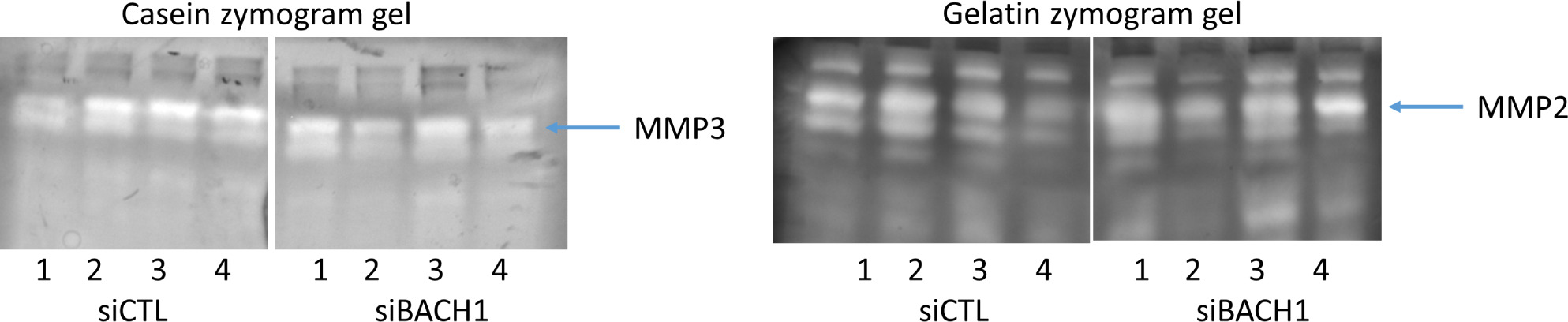
Effect of BACH1 silencing on MMP3 and MMP1. the MMP3 gel on the left (showing difference between siCLT and siBACH1) in the MMP-3 band and on the right the MMP-2 gel with no difference.

### C Gene nomenclature

ACSL6 (Acyl-CoA Synthetase Long Chain Family Member 6),

ALDH1A2 (Aldehyde Dehydrogenase 1 Family Member A2),

BACH1 (BTB Domain And CNC Homolog 1),

BBX (BBX High Mobility Group Box Domain Containing),

BHLHE40 (Basic Helix-Loop-Helix Family Member E40),

CBFB (Core-Binding Factor Subunit Beta),

CPT1C (Carnitine Palmitoyltransferase 1C),

CREM (cAMP Responsive Element Modulator),

ELF1 (E74 Like ETS Transcription Factor 1),

ELF4 (E74 Like ETS Transcription Factor 4),

ELF5 (E74 Like ETS Transcription Factor 5),

ENO1 (Enolase 1),

EGR2 (Early Growth Response 2),

ETS1 (ETS Proto-Oncogene 1, Transcription Factor),

ETV2 (ETS Variant Transcription Factor 2),

ETV3 (ETS Variant Transcription Factor 3),

ETV7 (ETS Variant Transcription Factor 7),

FLI1 (Fli-1 Proto-Oncogene, ETS Transcription Factor),

FOS (Fos Proto-Oncogene, AP-1 Transcription Factor Subunit),

FOSL1 (FOS Like 1, AP-1 Transcription Factor Subunit),

FOSL2 (FOS Like 2, AP-1 Transcription Factor Subunit),

FOXC2 (Forkhead Box C2),

HADHB (Hydroxyacyl-CoA Dehydrogenase Trifunctional Multienzyme Complex Subunit Beta), HIF1A (Hypoxia Inducible Factor 1 Subunit Alpha),

HIVEP1 (HIVEP Zinc Finger 1), HLX (H2.0 Like Homeobox), HOXA1 (Homeobox A1), HOXB2 (Homeobox B2), HOXB3 (Homeobox B3), HMOX1 (Heme Oxygenase 1),

IKZF1 (IKAROS Family Zinc Finger 1), IRF7 (Interferon Regulatory Factor 7),

JUNB (JunB Proto-Oncogene, AP-1 Transcription Factor Subunit), LINC01600 (Long Intergenic Non-Protein Coding RNA 1600), MECOM (MDS1 And EVI1 Complex Locus),

MITF (Melanogenesis Associated Transcription Factor),

MMD (Monocyte To Macrophage Differentiation-Associated), NF1 (Neurofibromin 1),

NFATC1 (Nuclear Factor Of Activated T Cells 1), NFE2L2 (NFE2 Like BZIP Transcription Factor 2), NKX2-1 (NK2 Homeobox 1),

NUDT15 (Nudix Hydrolase 15),

NR4A1 (Nuclear Receptor Subfamily 4 Group A Member 1),

PAX8 (Paired Box 8),

PKNOX1 (PBX/Knotted 1 Homeobox 1),

PLAGL1 (PLAG1 Like Zinc Finger 1),

RARG (Retinoic Acid Receptor Gamma),

RFX5 (Regulatory Factor X5),

RORC (RAR Related Orphan Receptor C),

RUNX1 (Runt Related Transcription Factor 1),

SLC25A26 (Solute Carrier Family 25 Member 26),

SLC35A5 (Solute Carrier Family 35 Member A5),

SMAD4 (SMAD Family Member 4),

STAT1 (Signal Transducer And Activator Of Transcription 1),

STRADB (STE20-Related Kinase Adaptor Beta),

SYNE2 (Spectrin Repeat Containing Nuclear Envelope Protein 2),

TAL1 (TAL BHLH Transcription Factor 1),

TBX4 (T-Box Transcription Factor 4),

TGIF1 (TGFB Induced Factor Homeobox 1),

TLX3 (T Cell Leukemia Homeobox 3),

TRIB3 (Tribbles Pseudokinase 3),

ZBTB1 (Zinc Finger And BTB Domain Containing 1),

ZBTB3 (Zinc Finger And BTB Domain Containing 3),

ZNF469 (Zinc Finger Protein 469).

## Bibliography

[1] Josef S Smolen, Daniel Aletaha, and Iain B McInnes. “Rheumatoid arthritis”. In: The Lancet 388.10055 (2016), pp. 2023–2038.

[2] Josef S. Smolen, et al. “Rheumatoid arthritis”. In: Nature Reviews Disease Primers 4 (2018).

[3] Yannis Alamanos and Alexandros A Drosos. “Epidemiology of adult rheumatoid arthritis”. In: Autoimmunity reviews 4.3 (2005), pp. 130–136.

[4] Josef S Smolen and Daniel Aletaha. “Rheumatoid arthritis therapy reappraisal: strategies, opportunities and challenges”. In: Nature Reviews Rheumatology 11.5 (2015), p. 276.

[5] Jackie L Nam et al. “Efficacy of biological disease-modifying antirheumatic drugs: a systematic literature review informing the 2013 update of the EULAR recommendations for the management of rheumatoid arthritis”. In: Annals of the rheumatic diseases 73.3 (2014), pp. 516–528.

[6] Adrià Aterido, et al. “A combined transcriptomic and genomic analysis identifies a gene signature associated with the response to anti-TNF therapy in rheumatoid arthritis”. In: Frontiers in immunology (2019), p. 1459.

[7] Ki-Jo Kim, et al. “Compendium of synovial signatures identifies pathologic characteristics for predicting treatment response in rheumatoid arthritis patients”. In: Clinical Immunology 202 (2019), pp. 1–10.

[8] Beatrix Bartok and Gary S Firestein. “Fibroblast-like synoviocytes: key effector cells in rheumatoid arthritis”. In: Immunological reviews 233.1 (2010), pp. 233–255.

[9] Fangqi Li, et al. “Nomenclature clarification: synovial fibroblasts and synovial mesenchymal stem cells”. In: Stem cell research & therapy 10 (2019), pp. 1–7.

[10] Melanie H Smith, et al. “Drivers of heterogeneity in synovial fibroblasts in rheumatoid arthritis”. In: Nature Immunology (2023), pp. 1–11.

[11] Richard I Ainsworth, et al. “Systems-biology analysis of rheumatoid arthritis fibroblast-like synoviocytes implicates cell line-specific transcription factor function”. In: Nature communications 13.1 (2022), pp. 1–11.

[12] Maryam Masoumi, et al. “Destructive roles of fibroblast-like synoviocytes in chronic inflammation and joint damage in rheumatoid arthritis”. In: Inflammation 44 (2021), pp. 466–479.

[13] TCA Tolboom, et al. “The invasiveness of fibroblast-like synoviocytes is of relevance for the rate of joint destruction in patients with rheumatoid arthritis and is a patient characteristic”. In: Arthritis Research & Therapy 7.1 (2005), pp. 1–1.

[14] Teresina Laragione et al. “The arthritis severity locus Cia5d is a novel genetic regulator of the invasive properties of synovial fibroblasts”. In: Arthritis & Rheumatism: Official Journal of the American College of Rheumatology 58.8 (2008), pp. 2296–2306.

[15] Yukinori Okada, et al. “Genetics of rheumatoid arthritis: 2018 status”. In: Annals of the rheumatic diseases 78.4 (2019), pp. 446–453.

[16] Eunji Ha, Sang-Cheol Bae, and Kwangwoo Kim. “Large-scale meta-analysis across East Asian and European populations updated genetic architecture and variant-driven biology of rheumatoid arthritis, identifying 11 novel susceptibility loci”. In: Annals of the rheumatic diseases 80.5 (2021), pp. 558–565.

[17] Alice M Walsh, et al. “Integrative genomic deconvolution of rheumatoid arthritis GWAS loci into gene and cell type associations”. In: Genome biology 17.1 (2016), pp. 1–16.

[18] Naouel Zerrouk, et al. “Identification of putative master regulators in rheumatoid arthritis synovial fibroblasts using gene expression data and network inference”. In: Scientific reports 10.1 (2020), pp. 1–13.

[19] Samuel A Lambert, et al. “The human transcription factors”. In: Cell 172.4 (2018), pp. 650–665.

[20] Fan Zhang, et al. “Defining inflammatory cell states in rheumatoid arthritis joint synovial tissues by integrating single-cell transcriptomics and mass cytometry”. In: Nature immunology 20.7 (2019), pp. 928–942.

[21] Fan Zhang, et al. “Deconstruction of rheumatoid arthritis synovium defines inflammatory subtypes”. In: Nature (2023), pp. 1–9.

[22] Oliver Stegle, Sarah A Teichmann, and John C Marioni. “Computational and analytical challenges in single-cell transcriptomics”. In: Nature Reviews Genetics 16.3 (2015), pp. 133–145.

[23] Aditya Pratapa, et al. “Benchmarking algorithms for gene regulatory network inference from single-cell transcriptomic data”. In: Nature methods 17.2 (2020), pp. 147–154.

[24] Pau Badia-i-Mompel, et al. “Gene regulatory network inference in the era of single-cell multiomics”. In: Nature Reviews Genetics (2023), pp. 1–16.

[25] Philipp Keyl, et al. “Single-cell gene regulatory network prediction by explainable AI”. In: Nucleic Acids Research 51.4 (2023), e20–e20.

[26] Seung Min Jung, Kyung-Su Park, and Ki-Jo Kim. “Deep phenotyping of synovial molecular signatures by integrative systems analysis in rheumatoid arthritis”. In: Rheumatology 60.7 (2021), pp. 3420–3431.

[27] Sungyong You, et al. “Identification of key regulators for the migration and invasion of rheumatoid synoviocytes through a systems approach”. In: Proceedings of the National Academy of Sciences 111.1 (2014), pp. 550–555.

[28] Suzi A Aleksander et al. “The Gene Ontology knowledgebase in 2023”. In: Genetics 224.1 (2023), iyad031.

[29] Minoru Kanehisa. “The KEGG database”. In: ‘In silico’simulation of biological processes: Novartis Foundation Symposium 247. Vol. 247. Wiley Online Library. 2002, pp. 91–103.

[30] David Croft, et al. “Reactome: a database of reactions, pathways and biological processes”. In: Nucleic acids research 39.suppl_1 (2010), pp. D691–D697.

[31] Matthew T Weirauch, et al. “Determination and inference of eukaryotic transcription factor sequence specificity”. In: Cell 158.6 (2014), pp. 1431–1443.

[32] Damian Szklarczyk, et al. “STRING v10: protein–protein interaction networks, integrated over the tree of life”. In: Nucleic acids research 43.D1 (2015), pp. D447–D452.

[33] Marieke Lydia Kuijjer, et al. “Estimating sample-specific regulatory networks”. In: Iscience 14 (2019), pp. 226–240.

[34] Matteo Manica, et al. “COSIFER: a Python package for the consensus inference of molecular interaction networks”. In: Bioinformatics (2020).

[35] L Shu, et al. “Mergeomics: Integrative Network Analysis of Omics Data”. In: (2017).

[36] Aurelien Pelissier, et al. “Cell-Specific Gene Networks and Drivers in Rheumatoid Arthritis Synovial Tissues”. In: bioRxiv (2023). pp. 2023–12.

[37] Masashi Miyazaki et al. “Tacrolimus and cyclosporine A inhibit human osteoclast formation via targeting the calcineurin-dependent NFAT pathway and an activation pathway for c-Jun or MITF in rheumatoid arthritis”. In: Clinical rheumatology 26.2 (2007), pp. 231–239.

[38] Tahir H Tahirov, et al. “Structural analyses of DNA recognition by the AML1/Runx-1 Runt domain and its allosteric control by CBF*β*”. In: Cell 104.5 (2001), pp. 755–767.

[39] Xinxia Sui, et al. “Association between IKZF1 related gene polymorphism, DNA methylation and rheumatoid arthritis in Han Chinese: A case-control study.” In: Authorea Preprints (2020).

[40] Shunichi Shiozawa and Ken Tsumiyama. “Pathogenesis of rheumatoid arthritis and c-Fos/AP-1”. In: Cell Cycle 8.10 (2009), pp. 1539–1543.

[41] Zhaolan Hu, et al. “The transcription factor RFX5 coordinates antigen-presenting function and resistance to nutrient stress in synovial macrophages”. In: Nature Metabolism 4.6 (2022), pp. 759– 774.

[42] Susan Hua and Thilani H Dias. “Hypoxia-inducible factor (HIF) as a target for novel therapies in rheumatoid arthritis”. In: Frontiers in pharmacology 7 (2016), p. 184.

[43] Gan Sun et al. “Loss of function mutation in ELF4 causes autoinflammatory and immunodeficiency disease in human”. In: Journal of Clinical Immunology (2022), pp. 1–13.

[44] Alfonso Martinez et al. “Role of SLC22A4, SLC22A5, and RUNX1 genes in rheumatoid arthritis.” In: The journal of Rheumatology 33.5 (2006), pp. 842–846.

[45] Marta E Alarcón-Riquelme. “Role of RUNX in autoimmune diseases linking rheumatoid arthritis, psoriasis and lupus”. In: Arthritis Res Ther 6.4 (2004), pp. 1–5.

[46] Zhiping Zhao et al. “Angiotensin II upregulates RANKL/NFATC1 expression in synovial cells from patients with rheumatoid arthritis through the ERK1/2 and JNK pathways”. In: Journal of Orthopaedic Surgery and Research 16.1 (2021), p. 297.

[47] Xuanan Li, et al. “Epigenetic regulation of NfatC1 transcription and osteoclastogenesis by nicotinamide phosphoribosyl transferase in the pathogenesis of arthritis”. In: Cell Death Discovery 5.1 (2019), p. 62.

[48] Susan E Sweeney, Maripat Corr, and Trevor B Kimbler. “Role of interferon regulatory factor 7 in serum-transfer arthritis: Regulation of interferon-*β* production”. In: Arthritis & Rheumatism 64.4 (2012), pp. 1046–1056.

[49] Kim Ohl, et al. “The transcription factor CREM drives an inflammatory phenotype of T cells in oligoarticular juvenile idiopathic arthritis”. In: Pediatric Rheumatology 16.1 (2018), pp. 1–9.

[50] Young-Mee Moon et al. “The Fos-related antigen 1–JUNB/activator protein 1 transcription complex, a downstream target of signal transducer and activator of transcription 3, induces T helper 17 differentiation and promotes experimental autoimmune arthritis”. In: Frontiers in immunology 8 (2017), p. 1793.

[51] Keiji Hirota, et al. “Autoimmune Th17 cells induced synovial stromal and innate lymphoid cell secretion of the cytokine GM-CSF to initiate and augment autoimmune arthritis”. In: Immunity 48.6 (2018), pp. 1220–1232.

[52] PV Kasperkovitz, et al. “Activation of the STAT1 pathway in rheumatoid arthritis”. In: Annals of the Rheumatic Diseases 63.3 (2004), pp. 233–239.

[53] Alistair LJ Symonds, et al. “Egr2 and 3 control inflammation, but maintain homeostasis, of PD-1high memory phenotype CD4 T cells”. In: Life Science Alliance 3.9 (2020).

[54] Evelyn P Murphy and Daniel Crean. “NR4A1-3 nuclear receptor activity and immune cell dysregulation in rheumatic diseases”. In: Frontiers in Medicine 9 (2022), p. 874182.

[55] Florian Renoux, et al. “The AP1 transcription factor Fosl2 promotes systemic autoimmunity and inflammation by repressing Treg development”. In: Cell reports 31.13 (2020).

[56] RW Kinne et al. “Synovial fibroblast-like cells strongly express jun-B and C-fos proto-oncogenes in rheumatoid-and osteoarthritis”. In: Scandinavian Journal of Rheumatology 24.sup101 (1995), pp. 121–125.

[57] Clément Guillou, et al. “Soluble alpha-enolase activates monocytes by CD14-dependent TLR4 signalling pathway and exhibits a dual function”. In: Scientific reports 6.1 (2016), p. 23796.

[58] Rizi Ai, et al. “Joint-specific DNA methylation and transcriptome signatures in rheumatoid arthritis identify distinct pathogenic processes”. In: Nature communications 7.1 (2016), p. 11849.

[59] Hironari Nishizawa, Mie Yamanaka, and Kazuhiko Igarashi. “Ferroptosis: regulation by competition between NRF2 and BACH1 and propagation of the death signal”. In: The FEBS Journal 290.7 (2023), pp. 1688–1704.

[60] Xiufeng Xie, et al. “BACH1-induced ferroptosis drives lymphatic metastasis by repressing the biosynthesis of monounsaturated fatty acids”. In: Cell Death & Disease 14.1 (2023), p. 48.

[61] Yoav Benjamini and Yosef Hochberg. “Controlling the false discovery rate: a practical and powerful approach to multiple testing”. In: Journal of the Royal statistical society: series B (Methodological) 57.1 (1995), pp. 289–300.

[62] Monali NandyMazumdar, et al. “BACH1, the master regulator of oxidative stress, has a dual effect on CFTR expression”. In: Biochemical Journal 478.20 (2021), pp. 3741–3756.

[63] Metello Innocenti. “New insights into the formation and the function of lamellipodia and ruffles in mesenchymal cell migration”. In: Cell Adhesion & Migration 12.5 (2018), pp. 401–416.

[64] Daniel Aletaha and Josef S Smolen. “Remission in rheumatoid arthritis: missing objectives by using inadequate DAS28 targets”. In: Nature Reviews Rheumatology 15.11 (2019), pp. 633–634.

[65] Andreas Kerschbaumer et al. “Efficacy of synthetic and biological DMARDs: a systematic literature review informing the 2022 update of the EULAR recommendations for the management of rheumatoid arthritis”. In: Annals of the rheumatic diseases 82.1 (2023), pp. 95–106.

[66] Marta F Bustamante, et al. “Fibroblast-like synoviocyte metabolism in the pathogenesis of rheumatoid arthritis”. In: Arthritis research & therapy 19.1 (2017), pp. 1–12.

[67] Rizi Ai, et al. “Comprehensive epigenetic landscape of rheumatoid arthritis fibroblast-like synoviocytes”. In: Nature communications 9.1 (2018), p. 1921.

[68] Carl Orr, et al. “Synovial tissue research: a state-of-the-art review”. In: Nature Reviews Rheumatology 13.8 (2017), pp. 463–475.

[69] Amanda Chan, et al. “The GTPase Rac regulates the proliferation and invasion of fibroblast-like synoviocytes from rheumatoid arthritis patients”. In: Molecular medicine 13.5 (2007), pp. 297–304.

[70] Teresina Laragione, Carolyn Harris, and Percio S Gulko. “TRPV2 suppresses Rac1 and RhoA activation and invasion in rheumatoid arthritis fibroblast-like synoviocytes”. In: International Immunopharmacology 70 (2019), pp. 268–273.

[71] Teresina Laragione, et al. “Huntingtin-interacting protein 1 (HIP1) regulates arthritis severity and synovial fibroblast invasiveness by altering PDGFR and Rac1 signalling”. In: Annals of the rheumatic diseases 77.11 (2018), pp. 1627–1635.

[72] Hironari Nishizawa et al. “Ferroptosis is controlled by the coordinated transcriptional regulation of glutathione and labile iron metabolism by the transcription factor BACH1”. In: Journal of Biological Chemistry 295.1 (2020), pp. 69–82.

[73] TA de Jong et al. “Altered lipid metabolism in synovial fibroblasts of individuals at risk of developing rheumatoid arthritis”. In: Journal of Autoimmunity 134 (2023), p. 102974.

[74] Joselyn Padilla and Jiyoung Lee. “A novel therapeutic target, BACH1, regulates cancer metabolism”. In: Cells 10.3 (2021), p. 634.

[75] Marta F Bustamante, et al. “Hexokinase 2 as a novel selective metabolic target for rheumatoid arthritis”. In: Annals of the rheumatic diseases 77.11 (2018), pp. 1636–1643.

[76] Marveh Rahmati, Mohammad Amin Moosavi, and Michael F McDermott. “ER stress: a therapeutic target in rheumatoid arthritis?” In: Trends in pharmacological sciences 39.7 (2018), pp. 610–623.

[77] Satoshi Wada, et al. “Bach1 inhibition suppresses osteoclastogenesis via reduction of the signaling via reactive oxygen species by reinforced Antioxidation”. In: Frontiers in Cell and Developmental Biology (2020), p. 740.

[78] Mark D Robinson and Alicia Oshlack. “A scaling normalization method for differential expression analysis of RNA-seq data”. In: Genome biology 11.3 (2010), pp. 1–9.

[79] Kimberly Glass, et al. “Passing messages between biological networks to refine predicted interactions”. In: PloS one 8.5 (2013), e64832.

[80] Abhijeet Rajendra Sonawane, et al. “Understanding tissue-specific gene regulation”. In: Cell reports 21.4 (2017), pp. 1077–1088.

[81] Camila M Lopes-Ramos, et al. “Sex differences in gene expression and regulatory networks across 29 human tissues”. In: Cell reports 31.12 (2020), p. 107795.

[82] Marouen Ben Guebila, et al. “GRAND: a database of gene regulatory network models across human conditions”. In: Nucleic Acids Research 50.D1 (2022), pp. D610–D621.

[83] Di Zhang, et al. “Identification of differentially expressed and methylated genes associated with rheumatoid arthritis based on network”. In: Autoimmunity 53.6 (2020), pp. 303–313.

[84] Nguyen Phuoc Long et al. “Efficacy of integrating a novel 16-gene biomarker panel and intelligence classifiers for differential diagnosis of rheumatoid arthritis and osteoarthritis”. In: Journal of clinical medicine 8.1 (2019), p. 50.

[85] Zhan-Chun Li, et al. “Functional annotation of rheumatoid arthritis and osteoarthritis associated genes by integrative genome-wide gene expression profiling analysis”. In: PloS one 9.2 (2014), e85784.

[86] Sumbul Afroz, et al. “A comprehensive gene expression meta-analysis identifies novel immune signatures in rheumatoid arthritis patients”. In: Frontiers in immunology 8 (2017), p. 74.

[87] Dmitry Rychkov, et al. “Cross-Tissue Transcriptomic Analysis Leveraging Machine Learning Approaches Identifies New Biomarkers for Rheumatoid Arthritis”. In: Frontiers in immunology 12 (2021), p. 2104.

[88] Jie Huang, et al. “Promising Therapeutic Targets for Treatment of Rheumatoid Arthritis”. In: Frontiers in Immunology 12 (2021), p. 2716.

[89] David S Wishart et al. “DrugBank 5.0: a major update to the DrugBank database for 2018”. In: Nucleic acids research 46.D1 (2018), pp. D1074–D1082.

[90] Kazuhiko Yamamoto, et al. “Genetics of rheumatoid arthritis in Asia—present and future”. In: Nature Reviews Rheumatology 11.6 (2015), pp. 375–379.

[91] Mulin Jun Li, et al. “GWASdb: a database for human genetic variants identified by genome-wide association studies”. In: Nucleic acids research 40.D1 (2012), pp. D1047–D1054.

[92] Janet Piñero et al. “DisGeNET: a discovery platform for the dynamical exploration of human diseases and their genes”. In: Database 2015 (2015).

[93] Sune Pletscher-Frankild, et al. “DISEASES: Text mining and data integration of disease–gene associations”. In: Methods 74 (2015), pp. 83–89.

[94] Allan Peter Davis et al. “The comparative toxicogenomics database: update 2019”. In: Nucleic acids research 47.D1 (2019), pp. D948–D954.

[95] Teresina Laragione, et al. “The cation channel Trpv2 is a new suppressor of arthritis severity, joint damage, and synovial fibroblast invasion”. In: Clinical immunology 158.2 (2015), pp. 183–192.

